# Systematic inference of mutation rates and spectra across the tree of life via a scalable read-based framework

**DOI:** 10.64898/2026.02.02.703326

**Authors:** Asaf Pinhasi, Keren Yizhak, Yosef E Maruvka

## Abstract

The rapid increase in available genome assemblies allows eukaryote-wide analyses of mutation rates and mutational spectra, yet whole-genome alignment remains a major computational bottleneck. We present CORAL, a scalable framework for inferring branch-specific substitutions without a centralized whole-genome alignment. CORAL fragments sister genomes into pseudo-reads, aligns them to an outgroup, and assigns substitutions by parsimony. CORAL achieved high concordance with three independent resources for both mutation rates and 96-category spectra. Applying CORAL to 5,090 species with calibrated divergence times, we generated the largest comparative atlas of mutation rates and spectra across animals, plants, fungi, and protists. Mutation rates vary by orders of magnitude and correlate with life-history traits such as lifespan and body weight. We find that mutation spectra are major determinants of each clade’s genomic trinucleotide composition and exhibit strong phylogenetic structure. We identified seven evolutionary mutational signatures, including two novel signatures and three previously observed only in cancer. Signature activities varied widely, and for several processes, tracked life-history covariates, suggesting distinct etiologies. Together, CORAL and this extensive atlas establish a powerful framework for comparative genomics, overcoming alignment bottlenecks to reveal the forces driving molecular evolution.

## Main

Mutation rates and mutational spectra are fundamental genomic traits of species ^1^. They determine how quickly genomes change and which sequence contexts are preferentially altered, thereby shaping the genomic material on which natural selection acts. Comparative genomics has been central to uncovering these patterns, primarily through whole-genome multiple sequence alignment (MSA), providing essential insights into genome evolution and species divergence.

In human cancer, “mutational signatures”, recurrent patterns in base-substitution spectra, have become a powerful framework for deconvolving underlying mutational processes ^2^. This framework has transformed our understanding of somatic mutagenesis and its biological and environmental drivers. In species evolution, however, comparable analyses have been applied far more narrowly, typically within limited taxonomic scopes or small numbers of genomes ^3,4^.

Recent advances in whole-genome sequencing have now produced thousands of high-quality assemblies across animals, plants, fungi, and protists ^5,6^, alongside extensive life-history and ecological trait databases ^7–9^. In principle, these resources should enable a global and systematic characterization of mutation rates and mutational signatures across the tree of life, and their association with underlying biological processes.

The tools that once powered comparative genomics have not kept pace with the scale and diversity of modern genome collections, leaving fundamental evolutionary questions underexplored. This limitation is largely methodological. Current comparative genomics pipelines are computationally intensive and constrain analyses to a fixed set of genomes. Reference-based “all versus one genome” strategies, such as Multiz ^10^, introduce a reference bias and cannot accommodate the full diversity of available assemblies. Reference-free alignments like those produced by Cactus ^11^ overcome these biases but remain difficult to scale to large numbers of species and require substantial computational resources to generate, update, or even query. As a result, existing alignment frameworks, while powerful for many comparative tasks, are cumbersome for mutation-rate and mutational spectrum analyses. They also lack the flexibility required for large-scale, species-specific substitution inference needed to analyze mutation rates and mutational signatures, particularly in scientific queries.

To address these challenges, we developed CORAL (**C**omparative **O**rthologous **R**ead-based **A**nalysis of **L**ineage Mutations), a lightweight and modular pipeline that infers lineage-specific substitutions by aligning pseudo-reads from sister taxa to an outgroup and assigning changes by parsimony. CORAL is designed for distributed computing, allowing for massive parallelization across thousands of species. We first evaluate key design choices (aligner, fragment length, and filtering) and benchmark CORAL’s substitution-rate estimates against established whole-genome alignment resources. We then apply CORAL across thousands of eukaryotic species triplets to generate a branch-resolved atlas of substitution rates and trinucleotide mutation spectra. Using this atlas, we (i) quantify cross-species variation in substitution rates and test associations with divergence time and life-history traits; (ii) derive and validate context-normalized spectra, examining how they relate to phylogeny and to genomic trinucleotide composition; and (iii) infer recurrent evolutionary mutational signatures and compare them to known somatic signatures while assessing their distribution across clades and their trait associations. Together, CORAL and the accompanying atlas provide a scalable framework for systematic comparative analyses of mutational processes across the tree of life.

## Results

### Overview of the CORAL tool

To enable fast and scalable genome comparison, CORAL (Figure 1) uses three-way genome comparison of closely related species. We first select a triplet of species in which two are more closely related (sister taxa), and the third serves as an outgroup. We then align the two sister taxa to the outgroup as follows: the sister taxa genomes are fragmented into short pseudo-reads (∼150 bp) with a 75 bp step size (∼2× tiling coverage), which are then aligned to the indexed outgroup genome using a local aligner (e.g., BWA-MEM ^12^; Methods). Pseudo-reads with low mapping quality or non-unique alignments are subsequently filtered out (Methods). We next traverse the outgroup genome and identify positions where reads from both sister taxa (species A and B) map. At each site, we compare the sister-taxon bases to the outgroup. A substitution is assigned to species A when it differs from the outgroup while species B matches it, and vice versa, indicating a change along the corresponding branch since divergence (Methods).

**Figure 1.**
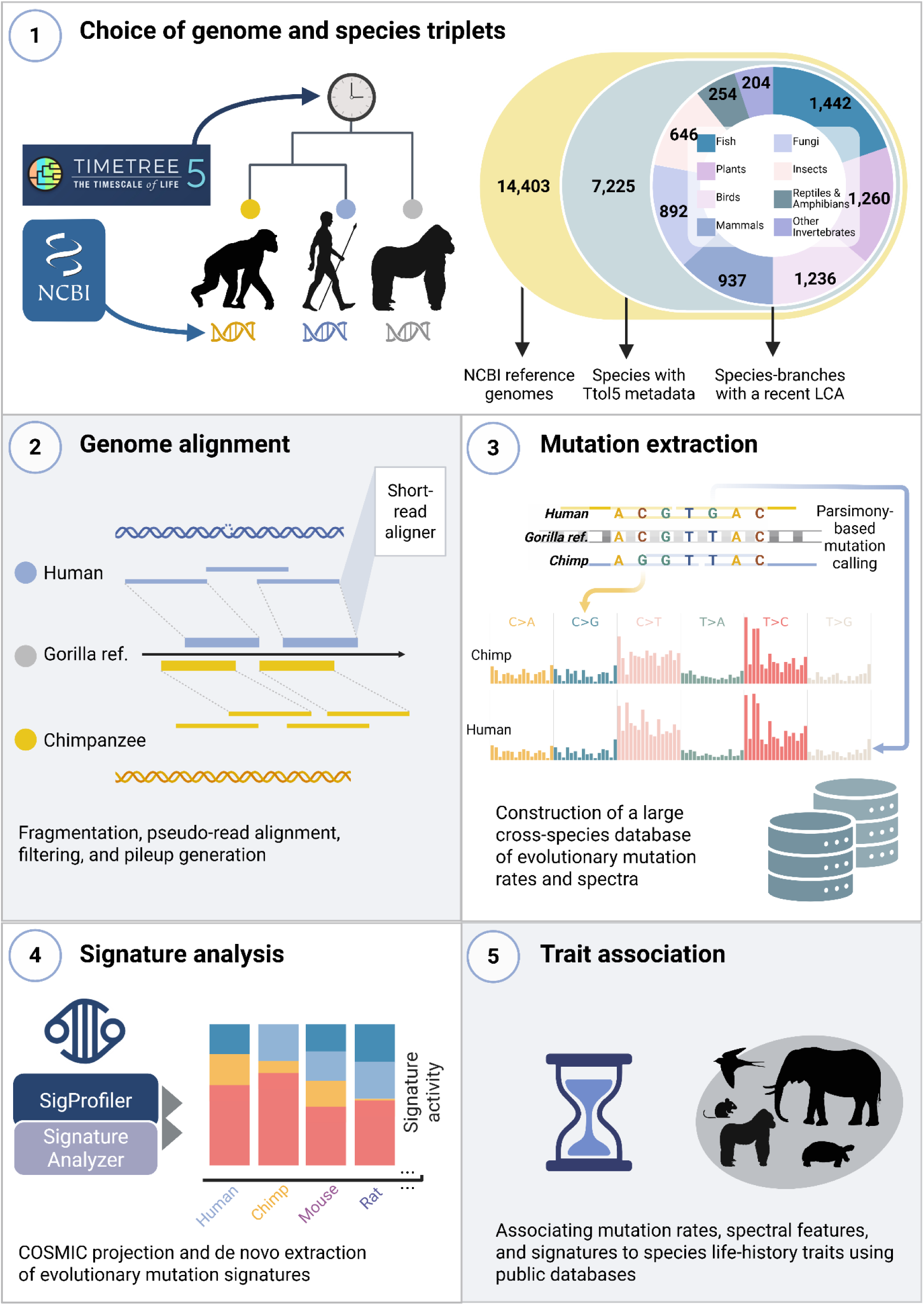
CORAL workflow and applications. **1,** Choice of genomes and species triplets: Species were selected based on the availability of NCBI reference genomes, curated TimeTree5 metadata, and phylogenetically close triplets, enabling outgroup-based mutation inference. **2,3 CORAL pipeline: 2.** Genome alignment: Each genome is fragmented and aligned using a short-read aligner to create local pseudo-read 3-way alignments. **3,** Mutation extraction: Mutations are inferred using a parsimony-based framework, generating a cross-species atlas of evolutionary mutation rates and spectra. **4,** Signature analysis: Mutation spectra were decomposed using COSMIC projections and de novo NMF-based extraction to identify conserved and species-specific evolutionary mutation signatures. **5,** Trait association: Mutation features were linked to species life-history traits obtained from public species life-history databases.

The method is highly efficient, as both the local read-mapping stage and the mutation-parsing process operate in near-linear time relative to genome size (Supplementary Note 1), allowing CORAL to be easily applied to thousands of species.

We benchmarked multiple aligners and fragment lengths across diverse species triplets (Supplementary Note 1). BWA-MEM ^12^ provided a consistently higher fraction of mapped reads (Figure S1a), while Minimap2 ^13^ was faster (Figure S1b). Fragment length had a similar trade-off: although the alignment fraction increased sharply from 50 bp to 150 bp, it plateaued beyond 150 bp while runtime - especially for large genomes - rose substantially (Figure S1c-d). Accordingly, we chose BWA-MEM with a read length of 150 bp as the optimal settings (Methods).

### CORAL mutation rates evaluation and dataset generation

We started by benchmarking CORAL against an established comparative resource. First, we estimated mutation rates by applying CORAL to a subset of species triplets that satisfied all inclusion requirements (Methods). For each lineage, we estimated mutation rates by calculating the number of substitutions per callable site (i.e., sites that were validly covered in all three species; Methods) and normalized it by sister taxa divergence time from TimeTree5 ^14^ (TToL5; Methods). We overlapped our dataset with the UCSC 241-way Zoonomia alignment and identified 76 species with matching CORAL and Zoonomia estimates.

We found a high correlation between the two estimates, with agreement improving as we increased minimum alignment coverage (Figure 2a): Pearson’s r increased from 0.95 for triplets with ≥ 25% alignment coverage to 0.99 for triplets with ≥ 70% coverage. Notably, our estimates were slightly lower than Zoonomia rates (Median difference 18%, Figure S2a), likely due to differences in the genomic regions analyzed. While we included all aligned regions, encompassing sites under purifying selection, Zoonomia focused on neutrally evolving sites. Despite these differences, the similarity increases when focusing only on species with high coverage, as demonstrated by the slope (Figure 2a).

**Figure 2.**
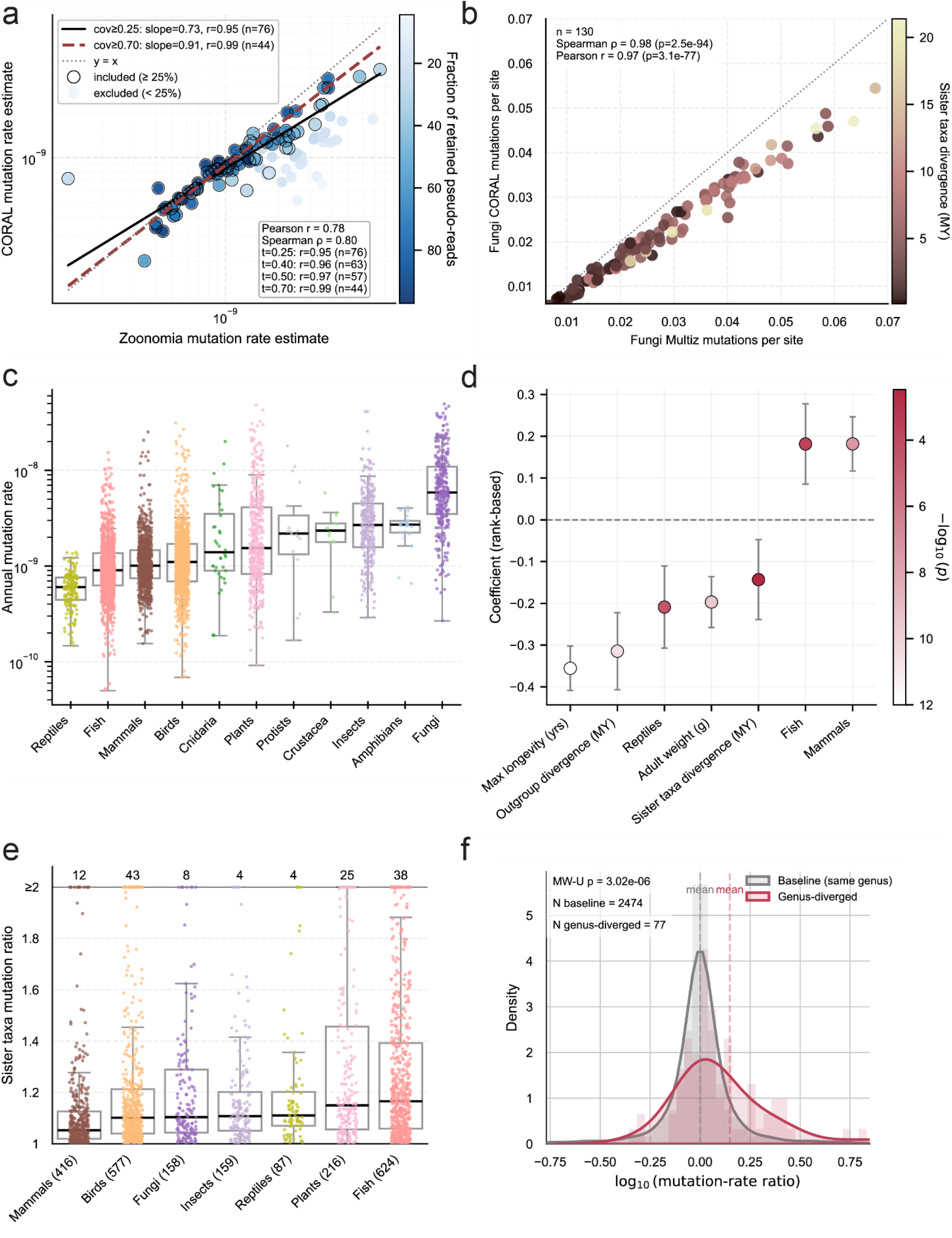
CORAL mutation-rate evaluation and mutation-rate variation across species. **a,** CORAL mutation-rates estimates compared with Zoonomia mutation-rates for 76 matched species-branches. Points are colored by the fraction of retained pseudo-reads; regression lines are shown for ≥25% and ≥70% alignment fraction thresholds. **b,** CORAL- vs Multiz-based mutations per site for chosen Fungi species-branches. Points are colored by sister taxa divergence time (MY). Two high-value outliers were excluded for visualization purposes. **c,** Annual mutation-rate distributions across clades. Boxplots show medians and interquartile ranges; points represent individual species-branches. For visualization purposes, only rates <5×10^-8^ are shown. **d,** Rank-based multivariate regression coefficients for predictors of mutation rate, including maximum longevity, adult body weight, sister-taxa and outgroup divergence, and clade identity (one-hot encoded). Error bars indicate standard deviation. **e,** Mutation-rate ratios between sister taxa across major clades. Points correspond to species pairs; numbers above plots indicate counts exceeding a twofold difference. **f,** Distribution of log10 mutation-rate ratios for same-genus triplets (grey) and genus-diverged triplets (red). Vertical dashed lines indicate the mean log10(ratio) for each group.

We next applied CORAL to all eukaryotic reference genomes in NCBI that met our inclusion criteria and had calibrated divergence times in TToL5 (Methods), yielding 5,363 species forming 3,472 sister-outgroup triplets with divergence times <60 million years (Tables S1-3). After removing triplets that have not passed the quality control (Methods), 3,460 triplets remained, representing 6,920 species-branches. After extraction, 49 additional branches with fewer than 1,000 inferred substitutions were excluded. The final dataset consists of 4,645 Metazoa, 1,260 Viridiplantae, 892 Fungi, and 74 Chromista/Protista species-branches, with some species appearing in multiple triplets, comprising 5,090 unique terminal species.

To further validate CORAL estimates beyond mammals, we analyzed a non-mammalian group by constructing three-way Multiz alignments for 130 fungal species (Figure 2b; Methods). We chose fungi, as their genome is relatively short (∼10-100 Mb) and they are evolutionarily distant from humans. In addition, fungal genomes are highly variable and show low alignment proportions (Figure S1g), making them a challenging test case while still allowing efficient construction of three-way MSAs. CORAL and Multiz estimates showed a remarkably high concordance (Spearman ρ = 0.98, *p* = 2.5×10^-94^, Pearson *r* = 0.97, *p* = 3.1×10^-77^). CORAL’s mutation rate estimates are modestly lower than Multiz (∼25% median difference; Figure S2a), but this bias is negligible relative to the three orders of magnitude variation across species (Figure 2a; Table S1). The bias reflects preferential alignment of conserved regions and is systematic, making it unlikely to confound associations with other features (Figure S2a). Conservation-related underestimation is inherent to multi-species genomic comparisons, affecting both multiple-sequence aligners (Multiz, Cactus) and triplet-based (CORAL) methods.

Importantly, each estimate reflects mutations that accumulated along the branch separating a species from its sister lineage. To avoid confusion with proper species-level traits, we refer to these estimates as species-branches. In addition, because taxonomic groups differ greatly in representation (e.g., many more animals than protists), comparison across traditional taxonomic ranks is problematic, especially for finer taxonomic resolutions. We therefore aggregated species into major, well-represented groups of interest, thus creating more interpretable categories for downstream analyses while retaining the broad evolutionary structure. Henceforth, we refer to these categories as “clades” (Table S1).

### Mutation rate variability

We estimated mutation rates for all species-branches (Table S1; Table S3). However, since low coverage may influence the estimated mutation rate, we focused here on species-branches with ≥25% retained pseudo-read fraction (n = 4,716; Methods; Supplementary Note 2). Rates varied both among and within taxonomic groups (Figure 2c). For instance, mammals showed a median rate of ∼1×10^-9^ mutations per bp per year, 1.5 times higher than reptiles (∼6×10^-10^). Birds displayed an even higher median (1.12×10^-9^), exceeded threefold by fungi (∼3.3×10^-9^). Within-group dispersion was also substantial: the 90th/10th-percentile ratio was 4.5 in mammals, 8.8 in insects, and nearly 18 in plants. Finer taxonomic structure (e.g., classes and orders) strongly contributed to this variability (Figure S3a).

Previous works have found that different biological and ecological factors correlate with mutation rates across species, including life-history traits such as body weight, life span, and litter size ^15–17^. However, given that many of these features also correlate with each other, it is not clear which is the main determinant of the correlation and what the contribution of each is. Our large dataset can enable multivariable analyses that overcome this intercorrelation. For that, we integrated life-history traits from AnAge ^7^ and AnimalTraits ^8^ with our atlas.

We started with an univariable analysis of the correlation between each trait and mutation rate (mean per taxon) (Tables S4; Figure S4a). As previously found, maximum longevity and adult body weight negatively correlated with annual mutation rates across clades. A few traits, such as birth weight and inter-litter interval, exhibited even stronger correlations, though in smaller subsets of species. Overall, the univariable results indicate that larger, longer-lived species with extended generation times, slower metabolic rates, and fewer offspring per year tend to accumulate fewer mutations. However, due to trait intercorrelation (Figure S4b), the contribution of each trait needs to be evaluated via multivariable analysis.

In mammals ^18^, generation time correlated with mutation rate more strongly than longevity did (Figure S4a). A partial rank correlation coefficient (PRCC; Methods) analysis including both traits identified generation time as the only significant independent predictor (Table S5), suggesting that it is the primary driver of yearly mutation accumulation between the two. An additional PRCC including maximum longevity, male maturity, and female maturity identified that longevity and male maturity were significant negative predictors, whereas female maturity was positive. This finding is consistent with the well-established male-biased contribution to germline mutation accumulation in mammals ^15,19,20^.

Divergence time has been widely shown to negatively correlate with mutation rate; that is, the estimated mutation rate tends to decrease as the divergence time between two species increases ^21–23^. Ho et al. ^22^ reviewed potential mechanisms to explain this, suggesting primarily that purifying selection eliminates mutations over longer timescales. We observed the same pattern across clades (Supplementary Note 3; Figure S5), with the exception of reptiles. Consequently, we included both sister-species divergence time and outgroup divergence time in our multivariable analysis.

To assess the joint contribution of key traits, we performed a rank-based multivariable regression including traits with the largest species coverage: (i) longevity, (ii) body weight, (iii) sister-taxa divergence, (iv) outgroup divergence, and (v) clade (birds as reference; Methods). Given the high variability of the estimates and the non-linear nature of these relationships, we used rank-based multivariable regression.

We found that longevity and body weight remained significant negative predictors of annual mutation rate in the multivariable model (Figure 2d, Table S5). A PRCC analysis (Table S5) confirmed that these associations reflect independent effects rather than shared variance, suggesting that despite their intercorrelation, each contributes independently to mutation-rate variation. The model further identified marked clade-level contributions: reptiles were associated with reduced mutation rates, whereas fish and mammals showed elevated rates relative to birds.

### Differences in mutation rates between sister taxa

To examine finer scale changes in mutation rates, we next asked how mutation rates vary between sister taxa. Most species pairs showed a balanced mutation-rate ratio (<1.3×; 2,645 of 3,435 pairs), yet a minority displayed pronounced differences, including 202 pairs with more than a twofold change (Figure 2e). This variability differs across clades, ranging from 91% stable sister taxa (<1.3 fold change) in mammals to ∼48% in Cnidaria. 10% of bird pairs and 7% of plant pairs exceeded a 2-fold difference, indicating faster mutation-rate evolution in these pairs.

We next investigated whether divergence into a new genus associates with mutation-rate shifts. We selected triplets where one sister taxon branched into a unique genus while the other remained in the same genus as the outgroup. In these triplets, the branching lineage exhibited a significantly higher mutation rate (p = 3.5×10^-5^, two-sided Wilcoxon signed-rank test; Figure S6). We further compared these triplets to a control set where all three species share the same genus. By analyzing the ratio of mutation rates between sister taxa, we found that rate asymmetry was significantly greater in branching lineages compared to stable, same-genus triplets (p = 3×10^-6^, two-sided Mann-Whitney test; Figure 2f). This demonstrates that a phenotypic change can be associated with an elevation in mutation rate.

### Mutation spectra

We next examined the variability in the rates of different substitution types. We began by analyzing transition-to-transversion (Ts:Tv) ratios and found that transitions exceeded transversions in nearly all species (Figure 3a; n = 6,770/6,871). Nonetheless, the Ts:Tv ratio varied substantially among clades. Medians ranged from 1.35 in Cnidaria to ∼2.2 in protists and mammals, peaking at ∼2.95 in fungi (two-sided Mann-Whitney test, p<10^-16^). Intra-class variation was also substantial; in fungi, the median Ts:Tv ratio differed over fourfold between Saccharomycetales and Ophiostomatales (Figure S7a,b). We validated the unexpectedly high fungal Ts:Tv ratio using three-way Multiz-based alignments (Figure S7d). Across 130 fungal branches, estimates were highly correlated between CORAL and Multiz (mean ratio was 2.93, 2.49, respectively) and significantly higher than other groups. Splitting the transitions and transversions into the six substitution classes (e.g., C>T, T>G) revealed even subtler clade-specific differences (Figure 3b).

**Figure 3.**
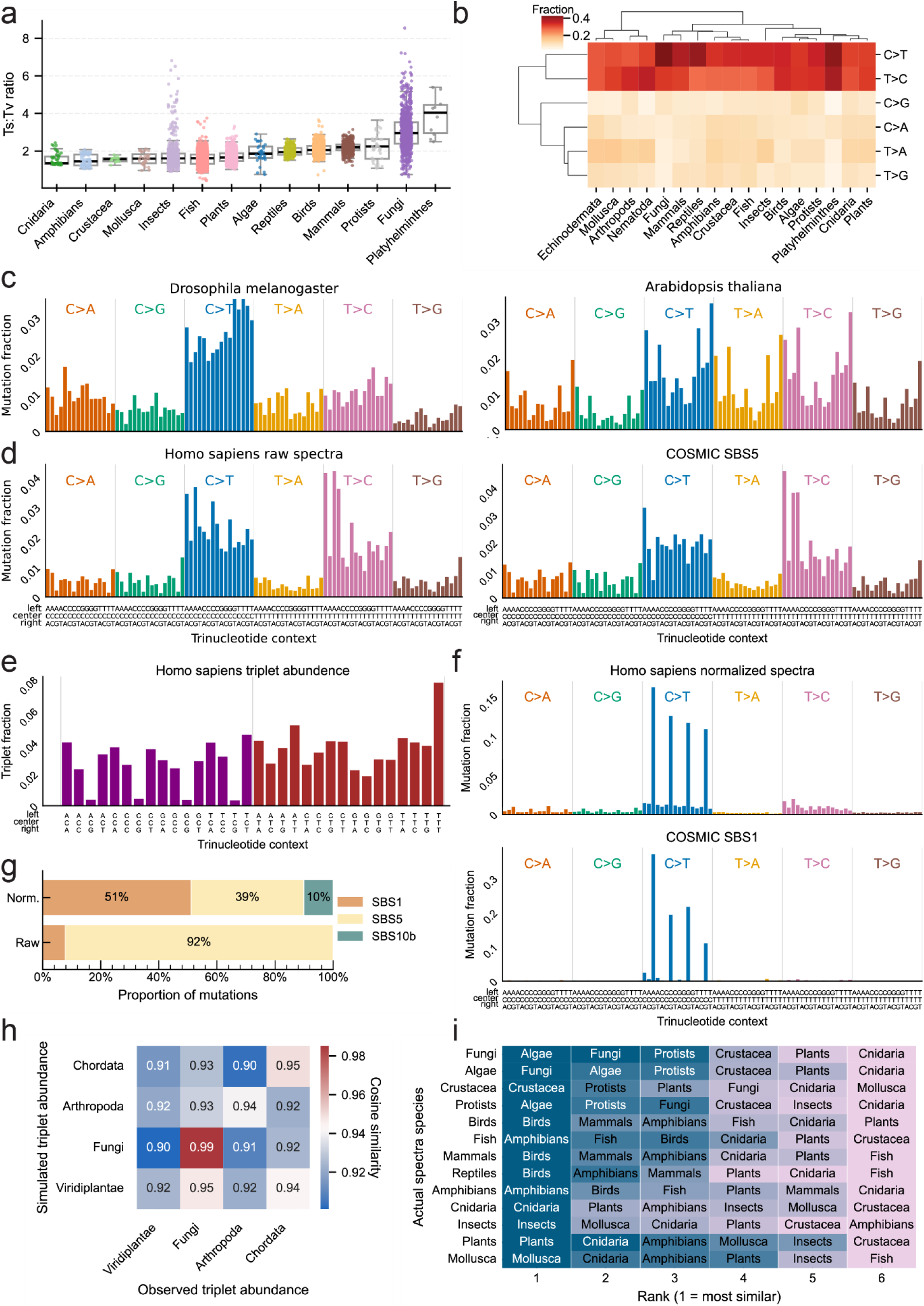
Mutation spectra composition and normalization. **a,** Transition/transversion (Ts:Tv) ratios across major clades, shown as boxplots; each point is a species-branch. **b,** Hierarchical clustering of six-class substitution spectra across clades. **c,** Example of raw 96-category mutation spectra for *Drosophila melanogaster* (left) and *Arabidopsis thaliana* (right). **d,** Raw human mutation spectrum (left) and COSMIC SBS5 spectrum (right). **e,** Trinucleotide abundance across the human genome. **f,** Normalized human mutation spectrum (top) and COSMIC SBS1 spectrum (bottom). **g,** Decomposition of raw and normalized human mutation spectra into COSMIC signatures using SigProfiler. **h,** Cosine similarity matrix comparing observed and spectrum-simulated trinucleotide abundances across major clades. **i,** Ranking heatmap of the top six groups whose spectrum-simulated 3-mer composition best predicts each clade’s observed composition (rows). Cell labels indicate the predicting group, and color encodes cosine similarity scaled within each row.

Finally, we examined the high-resolution 96-category mutation spectra, defined by substitution class and trinucleotide sequence context. These categories comprise all combinations of the six possible base substitution types (C>A, C>G, C>T, T>A, T>C, T>G), after collapsing reverse complements, with the sixteen possible 5′ and 3′ flanking nucleotide contexts ^24^. Spectra were derived from positions where flanking bases were conserved across all three species, and the central base was substituted in exactly one sister taxon (Methods).

The accuracy of CORAL inferred spectra was validated by comparison to three independent spectra resources: Multiz100 (vertebrates), Multiz-based alignments of 130 fungi species-branches, and msad213 (Supplementary Note 2). Across all datasets, CORAL-derived spectra showed strong concordance with the reference spectra, as quantified by cosine similarity. Median cosine similarity was highest for the fungal Multiz dataset (0.994), followed by msad213 (0.992) and Multiz100 (0.974). In all three comparisons, similarity between matching species was significantly higher than between non-matching species (Multiz100: mean 0.97 vs. 0.93, p = 2.5×10⁻⁷; fungi Multiz: 0.994 vs. 0.918, p = 2.47×10⁻⁸²; msad213: 0.987 vs. 0.954, p = 1.69×10⁻⁶; one-sided Mann-Whitney U tests, Supplementary Note 2). The slightly lower concordance observed in Multiz100 likely reflects reference bias, as all species are aligned to the human genome irrespective of evolutionary distance. In contrast to mutation rate estimates, spectral concordance showed no dependence on alignment coverage (Supplementary Note 2; Figure S2).

We found that the 96-category mutation counts differed between species (Figure 3c, d-left) and closely tracked their taxonomic classification (Figure S8a). This is expected regardless of the mutational process, since the genomic trinucleotide composition differs between species (Figure S8b). To account for these genomic differences, we normalized spectra by each genome’s trinucleotide frequencies (Figure 3e, Figure 3f-top; Methods). This correction removes 3-mer composition biases, thereby enabling direct comparison between species and yielding spectra that better reflect the underlying mutational processes.

### The human mutation spectrum and SBS1 and SBS5 signatures

Before comparing spectra across the tree of life, we examined the effect of normalization on the human trinucleotide mutation spectrum, which revealed an interesting pattern. The raw (non-normalized) human spectrum showed a high similarity to COSMIC SBS5 (cosine similarity = 0.95; Figure 3d, Figure S8c), indicating SBS5 as the most dominant continuous mutational process active throughout human evolution. However, after normalization, the human spectrum was most similar to the COSMIC SBS1 signature (cosine similarity = 0.95; Figure 3f, Figure S8d). Unsupervised COSMIC signature projection ^25^ supported this, yielding an inferred contribution of >90% from SBS5 prior to normalization, shifting to ∼50% from SBS1 and ∼40% from SBS5 after normalization (Figure 3g; Methods).

SBS1 and SBS5 are both clock-like signatures, correlated with human age and with each other ^26^. SBS1 is attributed to spontaneous deamination of 5-methylcytosine, whereas the etiology of SBS5 remains unknown despite its ubiquity in somatic tissues.

The emergence of SBS1 as the closest match after normalization led us to hypothesize that long-term SBS1 activity has imprinted itself in the human genome as altered triplet composition. Such imprinting is known to occur: the high C>T mutation rate at CpG sites, driven by methylation, leads over time to CpG depletion in the genome ^27,28^. Normalizing by triplet abundance effectively increases the relative rate of CpG mutations (in the normalized space), yielding a spectrum more similar to SBS1. In contrast, SBS5 does not strongly skew trinucleotide abundance, likely because its mutation profile is more balanced across contexts and therefore has a less conspicuous long-term impact on genomic composition. These observations prompted us to ask more generally to what extent mutation spectra shape genomic composition across taxa.

### Mutation spectra shape the genome content

The balance between the creation and removal of A/T and C/G bases is a major force in genome evolution, influencing GC content, genome stability, codon usage, and recombination rates ^29^. We found that while most taxa exhibited near-equilibrium ratios of T/A creation to removal (complemented by G/C removal to creation), several taxa deviate markedly from it (Figure S8e).

Mutations can alter the genomic content more subtly by changing 3-mer abundances through context-specific substitutions. Because CORAL provides both mutational events and triplet abundances per species, we could directly examine such composition shifts. We therefore asked whether SBS mutation spectra determine the observed triplet abundances, and how close species are to their expected steady state. Specifically, we quantified the deviation of each group’s observed 3-mer composition from the equilibrium predicted by its own spectrum. Using the mean normalized spectra of four groups - plants, fungi, insects, and chordates - we simulated long-term equilibrium compositions by iteratively applying the expected changes implied by each group’s spectrum until convergence (Methods).

Comparing these simulated steady states with observed compositions revealed striking similarity (mean cosine similarity ≈ 0.94; Figure 3h). Pairwise comparisons showed high correlations across the four tested groups, with each group’s spectrum most accurately predicting its own composition. We then moved to higher resolution and tested all our clades. At the clade level, simulated 3-mer profiles typically ranked among the top two most similar to observed profiles, except in reptiles (Figure 3i; Figure S8f), suggesting that reptiles are relatively far from their equilibrium. These results directly demonstrate, for the first time at such a large scale, that mutation spectra are major determinants of genomic 3-mer composition. This is evident by both the high overall similarity between simulated steady-state and observed compositions and the preferential ability of each clade’s spectrum to recapitulate its own genomic composition.

### The phylogenetic character of mutation spectra in evolution

Next, we examined the normalized mutation spectra, which reflect mutational processes while controlling for species-specific 3-mer abundance. We observed substantial variation across species (Figure 4a). For example, *Mus musculus* shows pronounced N[C>T]G (CpG) peaks reflecting strong 5-methylcytosine deamination, whereas *Drosophila melanogaster*, which lacks DNA methylation, shows no CpG-driven signal; instead, its spectrum is dominated by elevated C>T rates with moderate increases in C>A and T>C substitutions.

**Figure 4.**
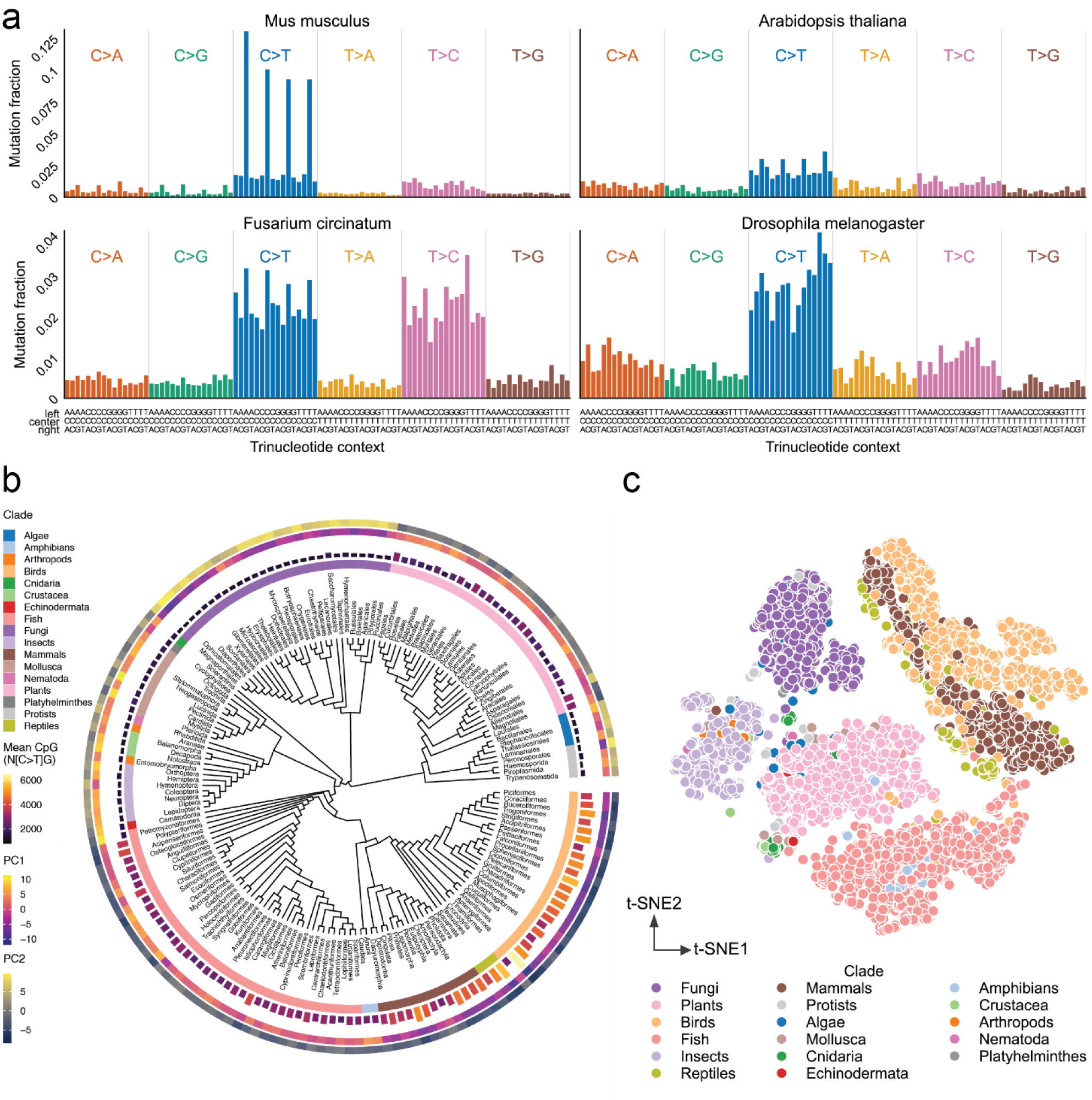
Phylogenetic structure of normalized mutation spectra. **a,** Example of normalized 96-category mutation spectra for representative species, including *Mus musculus* (top left), *Arabidopsis thaliana* (top right), *Fusarium circinatum* (bottom left), and *Drosophila melanogaster* (bottom right). **b,** Circular phylogeny annotated with clade identity (inner ring), mean CpG mutation levels (second ring), and principal component loadings derived from normalized mutation spectra (PC1, PC2). **c,** t-SNE embedding of normalized mutation spectra across species, with points colored by clade.

Similar to what was shown in mammals ^3,30^, we found that mutation spectra display clear phylogenetic structure across the whole eukaryotic tree-of-life (Figure 4b). Moreover, CpG mutation frequencies are high in chordates - especially mammals, birds, and reptiles - but are largely absent in invertebrates and fungi, with plants showing intermediate levels. Lampreys (Petromyzontiformes) are a notable exception among vertebrates, lacking a CpG peak, explained by their unique biology. Lampreys’ methylation system is considered intermediate between invertebrates and jawed vertebrates ^31^, undergoes substantial early-embryonic remodeling ^32^, and varies widely across individuals. These features are consistent with the transition in the vertebrate methylation system. Insects generally show low or absent CpG peaks, aligning with the well-documented variability of insect methylation and its complete loss in several major orders (e.g., Diptera, Hymenoptera) ^33^.

To extract unsupervised spectral patterns, we performed PCA on mean normalized spectra across orders (Methods). The first two components explained ∼45% of the variance and broadly separated major clades (Figure 4b, Figure S9). While PCA did not fully resolve all groups at the species level, t-SNE ^34^ (and UMAP ^35^) revealed clear clustering by clade (Figure 4c; Figure S10b; Methods), indicating that spectra are strongly associated with phylogeny but are not separable along two linear axes. Hierarchical clustering further supported this relationship (Figure S10c).

A subset of classes, consisting mainly of marine invertebrates including Gastropoda, Bivalvia, Anthozoa, Cephalopoda, and Echinoidea, clustered together in normalized spectral space despite being evolutionarily distant (Figure S10a-c). Their convergence may reflect shared aquatic environments, as well as differences in DNA-methylation patterns, including gene-body methylation and environmentally responsive methylation states.

To quantify how strongly mutation spectra reflect phylogenetic relationships, we performed Mantel tests. Both the random-permutation Mantel test and the phylogenetically constrained Phylo-Mantel test ^36^ (Methods) revealed a strong phylogenetic signal (p<<10^-4^), with observed correlations far exceeding all simulated permutations (Figure S10). Consistent with smaller-scale studies ^3,30^, these results show that mutation spectra are robust molecular traits encoding evolutionary relationships.

### Species mutational signatures

Inspired by cancer genomics, where somatic mutations are decomposed into characteristic signatures ^2^, we asked whether analogous “evolutionary signatures” could be identified, as was previously done on specific groups, e.g., apes ^3,4^.

To characterize evolutionary signatures, we first denormalized each species-branch trinucleotide spectra to the human trinucleotide abundance to enable a comparison with the human-based COSMIC signatures (Methods). We then extracted de novo signatures from the species denormalized spectra using unsupervised NMF (Methods) ^37^. A robust configuration yielded seven recurrent signatures across species (Figure 5a, Figure S11, Table S6). Of those, SBS1-like and SBS5-like were previously found in species evolution (Figure 5a) ^3^. Three were found in cancer development but not in species evolution: SBS30-like (likely related to base excision repair deficiency), SBS54-like (proposed germline variant contamination, or mismatch repair related signature), and SBS12-like (unknown etiology) ^2,24^. In addition, we found two novel signatures, SBSA and SBSB (Figure 5a; Figure S11). Most signatures, including SBS5-like, SBS30-like, SBSA, and SBS54-like, present a gradual increase across clades, whereas the other three (SBS1-like, SBSB, and SBS12-like) show a distinct activity between different clades. SBS1-like is mainly active in species with active DNA methylation mechanisms (Figure 5b,c), such as mammals and birds, while it is absent in insects and fungi, which lack this mechanism. This is aligned with our previous CpG-related observations in the spectra analysis section (Figure 4b). Interestingly, when comparing SBS1-like activity with methylation levels across species ^38^, a correlation between SBS1-like activity and methylation levels was observed only in birds (Figure 5d). SBS12-like was unique in its clade specificity, being largely restricted to fungi with minimal contribution in other lineages.

**Figure 5.**
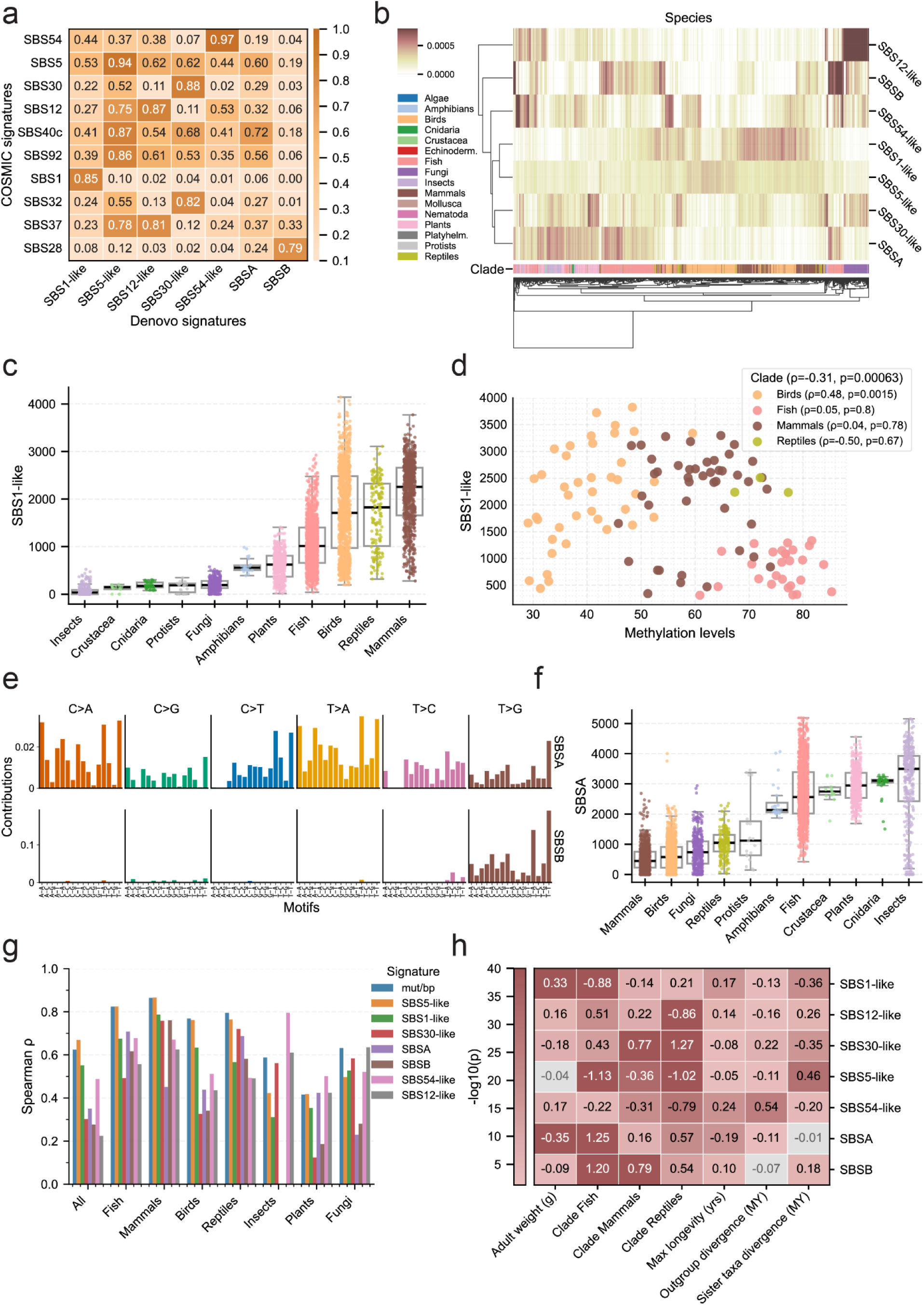
Evolutionary mutational signatures across species. **a,** Cosine similarity matrix comparing de novo evolutionary signatures with COSMIC SBS signatures. Rows correspond to COSMIC SBS signatures and columns to de novo signatures; values indicate cosine similarity. **b,** Heatmap of de novo signature activities across species. Rows correspond to species and columns to signatures; activities are normalized within each signature for visualization. Species are hierarchically clustered, with clade identity shown as an annotation bar. **c,** Distribution of SBS1-like signature activity across clades. Points represent species-branches; boxplots summarize per-clade distributions. **d,** Scatter plot of genomic methylation levels versus SBS1-like signature activity. Each point represents a species (mean SBS1-like activity) with available methylation level estimates; points are colored by clade. **e,** 96-category mutation spectra of the novel signatures SBSA and SBSB. Columns correspond to substitution classes and trinucleotide contexts; rows correspond to signatures. **f,** Distribution of SBSA signature activity across clades. Points represent species-branches; boxplots summarize per-clade distributions. **g,** Spearman correlation between signature and mutation burden per base vs sister-taxa divergence time. Bar height represents the Spearman correlation coefficient. **h,** Heatmap of regression coefficients from a multivariate rank-based model, fitting life-history and phylogenetic variables (rows) to signature activity (columns). Color denotes -log10(p-value); non-significant (>0.05) cells are greyed.

The novel SBSA, similar to SBS5-like, is a flat signature (Figure 5e), suggesting its etiology is likely related to general DNA damage or repair processes rather than a unique mechanism. Interestingly, this signature was found to be depleted in mammals, birds, and reptiles - amniotes with more sheltered germlines - while anamniotes, such as amphibians, fish, and insects, present high levels of activity (Figure 5f). This suggests that this signature may be related to reduced shielding from exogenous damage in externally developing gametes/embryos. However, this hypothesis does not explain the full picture, as fungi, for example, do not follow this trend.

The novel signature SBSB exhibits a unique, strong T>G bias (Figure 5e) and presents a low activity across most clades, except for a subset of fish having a larger contribution of SBSB (Figure S12). A finer comparison across fish orders (Figure S13) revealed an association between ambient temperature and SBSB activity. This suggests that a mechanism associated with adaptation to lower temperatures may be linked to the generation of this signature.

We next examined correlations between signature burden and sister-taxa divergence time (Figure 5g, Table S7; Methods). SBS5-like correlated with divergence time as strongly as, or in some clades more strongly than, total mutations per site, mirroring its time-dependent behavior in somatic tissues. In contrast, even in classes where all species-branches exhibited SBS1-like activity, SBS1-like correlations with divergence time were consistently weaker. This aligns with prior studies showing that SBS1 is not strictly time-dependent and suggests that SBS1 mutations may be more strongly subject to purification ^39^.

Finally, to disentangle life-history contributions to signature activity, we performed a rank-based multivariate regression, analogous to the mutation-rate model, using birds as a baseline (Figure 5h). Among all signatures, sister-taxa divergence time showed the strongest positive association with SBS5-like, and negative associations with SBS1-like and SBS30-like. SBS1-like also exhibited a strong positive correlation with adult body weight, whereas SBS5-like showed little dependence on either weight or longevity. These results are consistent with SBS5 accumulating linearly with time, whereas SBS1 reflects processes linked to cell division or saturation dynamics. Maximum longevity was most strongly associated with SBS1-like and SBS54-like. Altogether, these results suggest broadly conserved evolutionary signatures with clear parallels to known somatic processes.

### Further applications of CORAL

Up to here, we focused on genome-wide mutational features across species, namely mutation rates, spectra, and signatures, derived from simple three-way species comparisons. However, beyond these analyses, the distributed read-based design of CORAL can also support regional analysis of genomic changes. In Supplementary Note 4, we demonstrate how CORAL can be used to identify copy number changes analogous to CNV detection in cancer genomics. In addition, CORAL can support multi-species phylogenetic comparisons based on shared variable sites (Supplementary Note 4). Together, these results indicate CORAL’s potential to extend to capture finer-scale features of the evolutionary process and to illuminate mutational patterns along deeper evolutionary branches.

## Discussion

Comparative genomics increasingly relies on large-scale alignment datasets, yet the computational demands and rigidity of whole-genome multiple sequence alignments (MSAs) limit their use across taxonomically diverse species. Here, we presented CORAL, a distributed tool for inferring lineage-specific substitutions, explicitly designed to scale across parallel compute nodes. We benchmarked CORAL’s results against MSA results such as Multiz and Cactus (Zoonomia database) ^10,11,40^, and showed that CORAL recovers key evolutionary signals at a fraction of the computational cost of a classic MSA.

By applying CORAL to 6,920 species-branches, we have generated and made public the largest comparative atlas of lineage-specific mutation data assembled to date. We then used it to evaluate species mutation rates, characterize species mutation spectra, and identify new signatures active across the tree of life. We also showed that mutation rates vary by orders of magnitude across taxa and are strongly associated with life-history traits such as lifespan, adult body weight, and generation time, even after conducting a multivariable analysis.

When analyzing phylogenetic-based mutation rate estimates, it is important to distinguish between mutation rate - the rate at which genomic changes emerge - and substitution rate - the rate at which these changes become fixed in a population ^23,41^. While the fixation process may take tens of millions of years, the observable density of genomic changes starts to decline within a relatively short period (∼1 million years) and continues to do so for a long period due to purifying selection. Thus, the quantity measured here represents an intermediate state between the de novo mutation rate and the long-term substitution rate. This effect may explain the observed negative correlation between mutation rate and divergence time between sister taxa. We emphasize that capturing this intermediate quantity is a general characteristic of phylogenetic inference and not a limitation specific to CORAL.

Beyond absolute rates, we show that mutation spectra are robust, lineage-specific molecular traits with strong phylogenetic signals. We further demonstrate that these spectra are the primary architects of genomic composition. We found that the “steady-state” nucleotide frequencies predicted by a species’ mutation spectrum matched its actual genomic composition with high accuracy (cosine similarity ≈ 0.94). This implies that the nucleotide content of a genome (e.g., GC percentage) is largely determined by the neutral mutational biases active in its lineage over hundreds of millions of years of species evolution.

A particularly interesting finding is the distinct mutational profile of fungi. This clade shows a wide cross-species variation in mutation rates (12.2-fold 90th/10th percentile ratio) and the highest transition-to-transversion (Ts:Tv) ratios (∼3.0). The high transition-to-transversion ratio is likely due to the SBS12-like signature we identified, characterized by T>C mutations and strongly enriched in fungi.

The large scope of our research prevented us from diving deeper into lower taxonomic groups and conducting a class or order-based analysis. We should note that a finer analysis of smaller groups by themselves could identify more mutational signatures that are active in a small number of species or in underrepresented groups, such as subsets of plants, fungi, and insects.

Phylogeny-based estimates of mutation rates and spectra can be affected by alignment-driven selection bias: rapidly evolving, poorly conserved regions may fail to align to the reference genome and are therefore excluded, which can skew both rate and spectrum estimates. A second limitation is outgroup choice - when the outgroup/reference is evolutionarily distant, local alignment quality can degrade in some regions, potentially generating artifacts. These caveats apply broadly to phylogenetic alignment frameworks (e.g., Multiz and Cactus). However, CORAL may be more sensitive in this respect because it fixes the outgroup and treats it explicitly as the reference genome. Nonetheless, because Multiz is computationally intensive, it necessitates centralized alignments to a single reference (e.g., human) that is typically substantially more distant than CORAL’s outgroup, thereby drastically exacerbating this bias.

While focused on substitutions, CORAL can be extended to other genomic variations, for example, using positional data to detect copy-number variations (CNVs) and chromosomal asymmetries, as we demonstrated in *Amazona* birds. In addition, the framework could be generalized to reconstruct ancestral mutation spectra across larger phylogenies, including more than just triplets. As a distinct application, CORAL could offer a scalable scaffold for fine-grained MSA refinement, providing an efficient complement to full alignments for broad evolutionary questions.

Finally, given the continued reduction in sequencing costs and the vast expansion of sequencing across new species, it is expected that tens of thousands or even hundreds of thousands of genomes will be generated in the near future. CORAL holds the promise of enabling a scalable, simple way to analyze these large datasets.

## Methods

### CORAL’s alignment, quality control, and read filtration

Genomes of the two sister taxa (A and B) were fragmented into 150 bp pseudo-reads with a 75 bp step size (∼2× tiling coverage). Pseudo-reads were aligned independently to the indexed outgroup genome (O) using BWA-MEM (v1) ^12^ unless otherwise specified. Mitochondrial sequences and alignments to unplaced or alternative contigs (e.g., containing “Un”, “random”, “alt”, “fix”, or “hap”) were excluded.

Reads were included or filtered based on the following criteria:

i. High MAPQ (MAPQ=60) reads were retained;
ii. A pseudo-read with intermediate mapping quality (1<MAPQ<60) was retained if it overlapped on the outgroup genome, with one of the reads it overlapped with in the original genome (either upstream or downstream);
iii. Low MAPQ reads (MAPQ=0) were excluded.

This filtering procedure preferentially preserves both uniquely mapped reads and lower-MAPQ reads that participate in continuous local alignments, while eliminating isolated or unreliable mappings. The resulting filtered alignments were sorted by position on the outgroup and used to construct the three-species pileup from which substitutions were called.

### Three-species parsing and mutation assignment

Mutation calls were derived from a three-species pileup anchored on the outgroup genome. A site was considered callable only when both sister taxa (A and B) had at least one retained pseudo-read covering the position, and all reads within each species agreed on the nucleotide at that site. Sites at which all three species carried different nucleotides were excluded as non-callable.

To ensure unambiguous contextual assignment, mutations were evaluated only at sites where the flanking nucleotides (one base upstream and downstream) were identical across all three species. At such positions, a substitution was assigned to the lineage whose central base differed from the shared state of the other two. Specifically:

- If A differs while B and O match, the mutation is assigned to A.
- If B differs while A and O match, the mutation is assigned to B.

For example, for:

A: A-G-T

B: A-A-T

O: A-A-T

Mutation A[A>G]T is assigned to A.

Mutation calling was performed in a single linear scan of the outgroup genome, yielding counts for all 192 trinucleotide substitution types. These were subsequently collapsed by reverse-complement symmetry into the standard 96-category representation, in which the central base is restricted to C or T. Triplet abundances used for normalization were counted only from callable sites, and the 64 trinucleotide contexts were correspondingly collapsed into 32 reverse-complement categories.

### Multiz mutation calling and fungi three-way alignment construction

Mutation calling from Multiz alignments was performed analogously to CORAL mutation calling, and according to the logic described in *Three-Species Parsing and Mutation Assignment*. Only positions where all three species were aligned and identical in the immediately flanking nucleotides were considered.

For benchmarking in fungi, 65 species triplets with CORAL alignment coverage greater than 0.25 were randomly selected. For each triplet, a three-way multiple sequence alignment was constructed by first independently aligning each sister species to the outgroup using segalign ^42^, then merging both alignments into a three-species alignment using Multiz. Segalign and Multiz were both run with default parameters.

### Triplet selection and dataset construction

To construct species triplets for mutation inference, we applied the following selection and filtering procedure. First, all eukaryotic genomes in NCBI ^6^ with assembly at least at the scaffold level were selected. A time-calibrated tree for all species was constructed using TToL5 ^14^, and species lacking calibrated divergence times were removed. For each remaining species, the closest relative by divergence time was identified as its sister taxon; when multiple equally close species were available, the sister taxa were chosen randomly.

For each sister pair (A, B), an outgroup (O) was selected by moving to the next ancestral node and choosing a descendant lineage whose divergence from the sister pair exceeded the sister divergence. Triplets in which the sister taxa diverged from the outgroup over 60 MYA were excluded. In addition, triplets were required to satisfy one of two separation constraints to ensure correct evolutionary orientation: dist(A, B) × 1.3 < dist(A/B, O) or dist(A, B) + 3 MYA < dist(A/B, O). Triplets violating this criterion were discarded. Additional exclusions were applied to triplets containing species with missing or corrupted FASTA assemblies and genomes whose alignment required more than three days.

After filtering, 3,460 triplets remained from the set of candidates diverging <60 MYA. Each triplet contributed two lineage branches (one per sister taxon), yielding 6,920 branches before downstream branch-level filtering. Notably, species may participate in multiple triplets, acting as a sister taxon in some and as an outgroup in others.

### Branch-level filtration criteria

Filtration was applied at the branch level after mutation extraction by two criteria:

1. Minimum mutation count:
Branches with < 1,000 assigned substitutions were excluded.
2. Minimum alignment coverage:
For de novo signature extraction, mutation rate and signature burden association to traits (Figure 2b-c, Figure 5) - only branches with ≥ 25% aligned pseudo-reads were retained. This threshold follows the benchmark indicating high accuracy for mutation rate estimates above this coverage level (see Figure 2a).

### Mutation-rate estimation

Branch-specific mutation rates were computed as follows:

To estimate the mutation rate, we first calculated the proportion of callable sites containing a mutation. We then normalized this fraction by the divergence time between the two sister taxa as computed by TToL5. Mathematically:

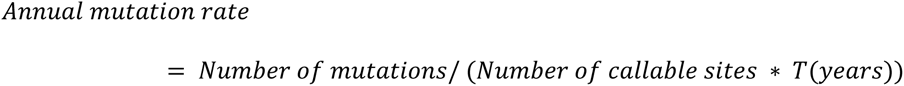

For Zoonomia mutation rates, we used neutral substitutions per site estimated from the UCSC phylogenetic tree. We converted these estimates to per-year rates by dividing them by the corresponding branch lengths *T*(*years*).

### Integration of life-history and ecological traits

Species traits were compiled from multiple curated databases, including AnAge, AnimalTraits, and Pacifici et al. (generation length and longevity) ^7,8,18^. The merged dataset includes morphological, reproductive, lifespan, and metabolic data. Trait variables were matched to species by name, and only branches with non-missing values for a given trait were included in each analysis. When appropriate, multiple branches belonging to the same species were averaged to produce species-level values.

### Univariate trait-mutation rate associations

For each life-history trait, we first aggregated branch-specific estimates to the species level by averaging the annual mutation rate across all branches belonging to the same species. Univariate Spearman correlations between species-level traits and species-level mutation rates were then computed in two settings: (i) across all species, and (ii) within each clade. Clade-level correlations were reported only when at least 25 species contributed data for that trait. Results are summarized in Table S4.

### Multivariate and PRCC analysis

Multivariate trait association with mutational features was performed in rank space rather than using the raw feature values. The target feature (either mutation rate or signature fraction) was modeled using ordinary least squares (OLS) regression with HC3 heteroscedasticity-robust standard errors, implemented with Python’s *statsmodels* package ^43^.

The predictor set included:

1. Maximum longevity and adult body weight (from AnAge ^7^);
2. Sister-taxa and outgroup divergence times (TToL5 ^14^);
3. One-hot encoded categorical variables for major clades (Reptiles, Mammals, and Fish), with Birds serving as the reference category to avoid perfect collinearity.

The mutation rate partial rank correlation coefficient (PRCC) analysis was conducted on the following groups of traits:

- G1: Adult weight, Maximum longevity, Sister-taxa divergence time, Outgroup divergence time
- G2: Female maturity, Male maturity, Maximum longevity, Sister-taxa divergence time, Outgroup divergence time
- G3: Calculated generation length, Maximum longevity difference, Sister-taxa divergence time, Outgroup divergence time
- G4: Birth weight, Adult weight, Sister-taxa divergence time, Outgroup divergence time

PRCC was performed with the *pingouin* Python package ^44^. The method estimates the partial Spearman correlation of each predictor with the target variable while controlling for all other predictors. PRCC dissects the unique contribution of the predictors to the target, independent of the predictors’ shared variance.

### Sister-taxon rate ratios and genus-branching analyses

To quantify mutation-rate asymmetry between sister taxa, we computed the sister-taxon ratio:

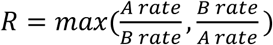

which equals 1 for perfectly balanced rates and increases with asymmetry.

To test whether branching into a new genus is associated with elevated mutation rates, we classified triplets into:

1. Genus-conserved triplets: all three species share the same genus.
2. Genus-diverged triplets: one sister taxon belongs to a different genus, while the other sister and the outgroup retain the same genus.

Two statistical tests were performed:

- Mann–Whitney U test comparing genus-conserved vs genus-diverged ratios.
- Paired Wilcoxon test comparing the diverging taxon to the conserved taxon species-branches within each genus-diverged triplet.

Both were performed on all species-branches without filtration.

### Spectra normalization and scaling

To account for genomic compositional differences, mutation counts in each context were normalized by the abundance of the corresponding trinucleotide in that species’ genome. For example, the mutation count for A[T>G]C was divided by the total number of ATC occurrences in the species’ callable genome (identical to the outgroup’s abundance). This results in normalized mutation frequencies for each of the 96 categories.

To allow standardized comparison of spectra across species with different genome sizes, mutation rates, and divergence times, each normalized spectrum was scaled to the same number. While normalizing to one would be standard, we normalized to 10,000 and rounded to the closest integer in order to fit the requirements of signature inference. This rescaling preserves the relative proportions of mutation types while placing all spectra on a common scale.

For comparisons involving COSMIC signatures, a human-denormalized spectrum was generated by multiplying each species’ normalized spectrum by the human trinucleotide abundances. For humans, this procedure reconstructs the original non-normalized spectrum; for other species, it yields spectra adjusted to human sequence composition, enabling direct signature comparisons on a shared background.

### Simulated equilibrium of triplet abundance

To evaluate to what extent mutation spectra explain the observed genomic 3-mer abundances, we simulated the expected steady-state triplet composition for each major group under its own mean normalized mutation spectrum.

Starting from a uniform triplet distribution, we performed iterative updates of the triplet-abundance vector. At each iteration:

1. The group’s normalized mutation spectrum was multiplied by the current 3-mer frequencies to obtain expected mutation counts under that composition.
2. These inferred mutation counts were used to compute the expected net change in each 3-mer class (creation minus removal).
3. The triplet-abundance vector was updated using a small learning rate (default 𝜂 = 1).
4. The process was repeated until the absolute sum of the update vector fell below a predefined convergence threshold (default 𝜀 = 10^−10^).

This procedure yields the predicted equilibrium triplet distribution implied solely by the species’ mutation spectrum. After convergence, simulated triplet abundances were compared to the observed genomic abundances using cosine similarity to assess how well mutational processes alone recapitulate lineage-specific 3-mer composition.

### Dimensionality reduction

We performed PCA on grouped mean spectra at several taxonomic levels (e.g., order, genus, taxa), using the 96 context-normalized mutation spectra as input. Group-level spectra were computed by averaging the normalized spectra of all species-branches in the group.

At the species–branch level, t-SNE ^34^ (default sklearn) was applied to three representations: raw 96-category spectra, context-normalized spectra, and genome 3-mer abundance profiles. UMAP^35^ (5 nearest neighbors, 0.5 min distance) was applied to context-normalized spectra. These embeddings were used to visualize clustering by taxonomic groups.

### Spectra phylogenetic relation quantification with Mantel test

To quantify how strongly mutation spectra reflect phylogeny, we compared a pairwise spectral distance matrix computed from species-level normalized spectra with the distance matrix derived from the time-calibrated phylogeny. A Mantel test with 10,000 permutations was used to estimate the correlation between these distances. In addition, a phylogenetically constrained Phylo-Mantel analysis was performed, in which permutations were restricted by phylogenetic proximity, providing a more conservative test of phylogenetic signal in the mutation spectra. Phylo-Mantel was performed using Evolqg’s PhyloMantel package in R ^36^.

### Signature inference and projection

De novo evolutionary signatures were inferred using the ARD-NMF implementation in SignatureAnalyzer ^37^ with phi=6 and COSMIC v3 as the reference signature set. SignatureAnalyzer automatically selects the number of inferred signatures. Human denormalized spectra of species-branches with over 25% pseudo-read alignment were provided as input.

To project known COSMIC signatures onto normalized and pre-normalized human spectra, we used SigprofilerAssignment (COSMIC v3.4) ^25^. Decomposition and de novo refitting were disabled to ensure direct projection of fixed reference signatures.

### Correlation with genomic methylation

Where species-level methylation estimates were available ^38^, Pearson and Spearman correlations were computed between genomic methylation fraction and SBS1-like activity. Analyses were performed separately within taxonomic groups (e.g., birds, fish) that had at least three species with methylation data.

### Time dependence of signature accumulation

To evaluate whether signatures accumulate with evolutionary time, we correlated each species’ signature burden (normalized activity × total mutation number) with sister-taxa divergence time. Correlations were computed separately for major clades.

## Data availability

Species reference genomes were downloaded from the NCBI Genomes Database 6 using accession numbers listed in Table S8. Divergence time estimates were obtained from TToL5 14 via the web portal and are provided in Table S1. The Multiz100 multiple alignment and Zoonomia substitution data were downloaded from the UCSC Genome Browser (https://hgdownload.soe.ucsc.edu/goldenPath/hg38/multiz100way/ and https://hgdownload.cse.ucsc.edu/goldenpath/hg38/cactus241way/, respectively). Msad213 mammalian spectra information 3, life-history datasets including AnAge 7, AnimalTraits 8, GL dataset (generation time estimates) 18, and cross-species methylation measurements 38, were obtained from the corresponding publications, and are included in Table S1. The mutation spectra dataset generated in this study, alongside different traits and genomic features, is available in Table S1 and in [CORAL_Data](https://drive.google.com/drive/folders/1Ydckj3TOsqvkJtDYfTY11BhOzw3tkYVm?usp=sharing). The full output data from CORAL, including positional mutation calling and alignment files, can also be downloaded via [CORAL_Data](https://drive.google.com/drive/folders/1Ydckj3TOsqvkJtDYfTY11BhOzw3tkYVm?usp=sharing).

## Code availability

The CORAL source code is publicly available at https://github.com/MaruvkaLab/CORAL. Scripts and data files used for data processing, analysis, and figure generation are provided in <u>CORAL_Data</u>.

## Acknowledgments

YEM was supported by the ISF grant number (2794/21), AP was supported by the Miriam and Aaron Gutwirth Memorial Scholarship. We would like to thank the Maruvka lab members for helpful discussions and support. Figure 1 was created with BioRender.com using a paid license.

## Supplementary Information

### Supplementary Note 1

#### CORAL performance in different settings

Genomic fragments from different species align with their outgroups in varying proportions due to genomic divergence and variation in reference genome completeness. Although some mutational features can be reliably inferred from a relatively small portion of the genome, others may be biased under low coverage. In particular, when genome coverage is low, retained pseudo-reads tend to originate primarily from conserved regions, which can lead to systematic underestimation of mutation rates. This issue affects all alignment-based genome comparison methods but warrants special attention here, as aligners differ in sensitivity.

We therefore evaluated the performance of several short-read aligners on a set of species triplets spanning diverse phylogenetic groups. Ten triplets with diverse genome sizes were selected, and CORAL was run using BWA-MEM^1^, Minimap2^2^, and BBMap^3^, each tested across multiple fragment lengths (Methods). BBMap exhibited unstable CPU usage and high memory demands, leading to unexpected pipeline failures, and was therefore excluded from further analysis.

BWA-MEM was more sensitive than Minimap2 in pseudo-read mapping, retaining a higher proportion of pseudo-reads aligned to the outgroup across nearly all tested triplets (Figure S1a). This advantage was most pronounced in species with a low overall fraction of retained pseudo-reads. In contrast, Minimap2 was substantially faster than BWA-MEM across fragment lengths and genome sizes (Figure S1b). Prioritizing sensitivity over runtime, and given that both aligners scale approximately linearly with genome size and can align large genomes within reasonable timeframes (<72 hours on a single CPU), we selected BWA-MEM for database generation and downstream analyses. The more recent BWA-MEM2 further improves runtime without altering alignment behavior, supporting this choice^4^.

As shown in Figure S1c,d, both the fraction of retained pseudo-reads and alignment time increase with fragment length. However, for BWA-MEM, while the alignment fraction rises steeply between 50 bp and 150 bp, it plateaus between 150 bp and 200 bp. On the contrary, alignment time - especially for large genomes - increases sharply over this range. Based on this trade-off, we selected 150 bp as the optimal fragment length for downstream analyses.

CORAL implementation natively supports all three aligners tested here, and users may also supply a custom aligner command, provided that standard input (FASTA/FASTQ) and output (BAM/SAM)^5^ formats are maintained. This ensures flexibility for future improvements and adaptation of alignment strategies to the evolutionary context.

Across all species, the fraction of retained pseudo-reads passing quality control (Methods) varied substantially (Figure S1e). This alignment fraction was strongly influenced by divergence time from the outgroup (Figure S1f) and by phylogenetic group (Figure S1g; Figure S3b), with additional contributions likely arising from genome assembly completeness and sequencing quality. Alignment fractions were generally higher in well-characterized clades such as Mammalia, Aves, Reptilia, and Actinopteri compared with most insects, plants, and fungi. Nonetheless, alignment-related biases are likely smaller than those associated with reference-based multiple sequence alignments, which typically retain only a limited subset of highly conserved regions relative to the reference genome.

Two notable deviations from these trends were observed. First, a subset of bird species overrepresented by parrots (*Psittaciformes*) showed relatively low fractions of retained pseudo-reads, despite having closely related outgroups (Figure S3c). Parrots are known to have unusually frequent events of chromosome fusions and fissions, as well as a high proportion of transposable elements compared to other birds ^6^. Second, reptiles showed the opposite pattern, maintaining high alignment fractions despite having the longest outgroup divergence times among all clades (Figure S3b). Specifically, Lacertidae (wall lizards) exhibited exceptionally high alignment (>55%), even though most diverged from their outgroups over 40 MYA, suggesting long-term genomic stability.

Importantly, alignment runtime scaled well with the target genome sizes (Figure S1h). Doubling of genome size increases runtime by roughly 2.5-fold, yielding a high Pearson correlation (r=0.78) between the total sister taxa genome sizes and alignment time. This scaling enables CORAL to align even the largest genomes within hours, supporting its broad applicability (Table S2).

**Figure S1.**
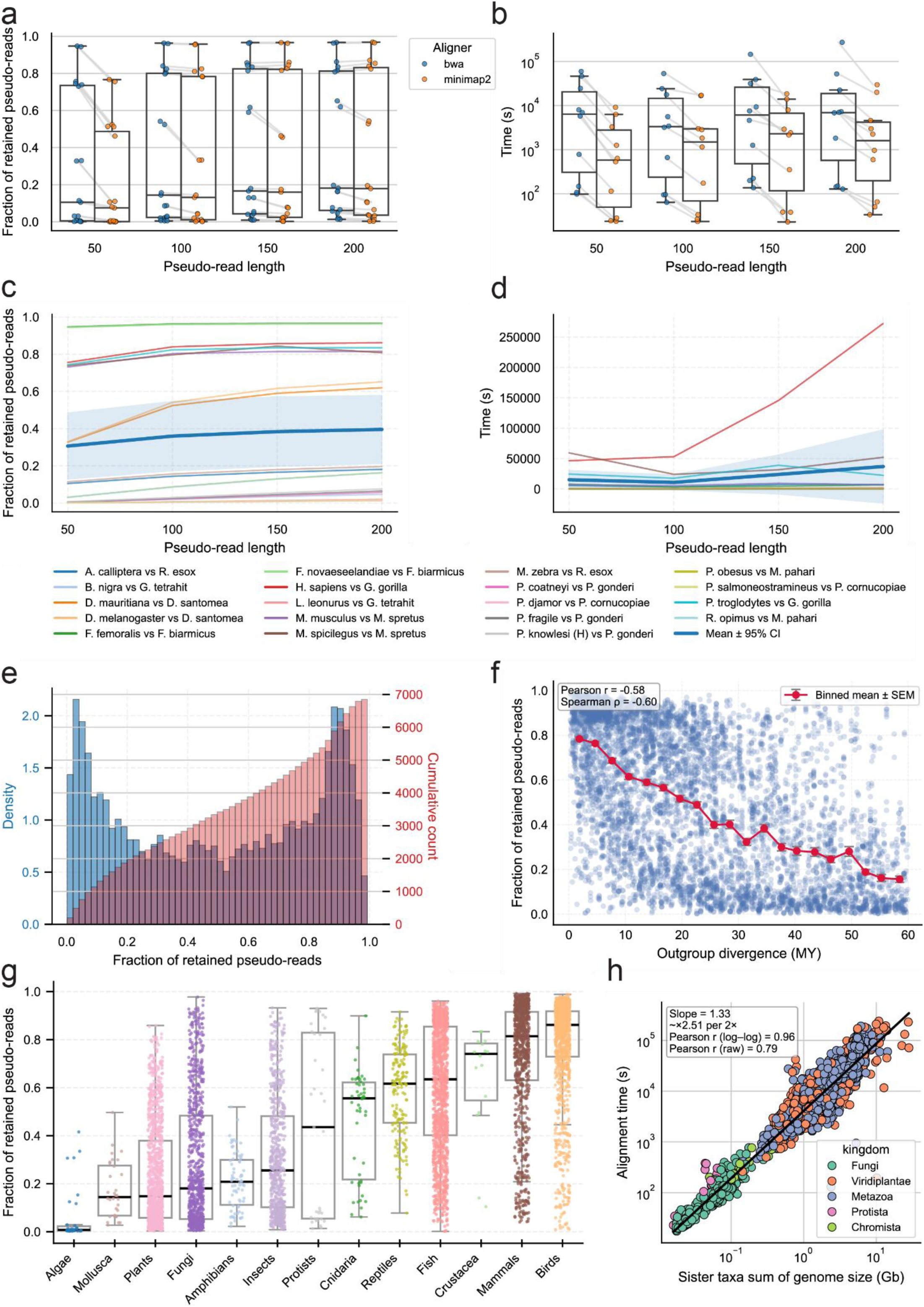
Alignment performance and coverage across species and aligners. **a,** Fraction of pseudo-reads retained after filtering for BWA-MEM (blue) and Minimap2 (orange) across pseudo-read lengths. Boxplots summarize selected species triplets; points represent species-branches, with lines connecting the same branches across aligners. **b,** Alignment runtime for BWA-MEM and Minimap2 across pseudo-read lengths, plotted as in **a**. **c,** Fraction of retained pseudo-reads versus pseudo-read length for selected species triplets using BWA-MEM. Thin lines indicate individual triplets; the thick line shows the mean ±95% confidence interval. **d,** Alignment runtime versus pseudo-read length for selected species triplets using BWA-MEM, plotted as in **c**. **e,** Distribution of the fraction of retained pseudo-reads across all analyzed species-branches, shown as a density histogram (blue) with cumulative count (red). **f,** Fraction of retained pseudo-reads versus outgroup divergence time (MY). Points represent species-branches; the red line shows the binned mean ± SEM. **g,** Fraction of retained pseudo-reads across clades. Boxplots summarize per-clade distributions with species-branch points overlaid. **h,** Alignment runtime versus the sum of sister-taxa genome sizes (Gb). Points represent species triplets, colored by kingdom; the fitted line indicates the scaling trend.

### Supplementary Note 2

#### External validation of CORAL spectra and species-branch filtration

To further validate our spectra, we performed three comparisons: to spectra extracted from the Multiz100 alignment using the same three-way approach (Methods), to spectra from *msad213* ^7^, and to spectra derived from three-way Multiz alignments in fungi we generated. For each species, we computed the cosine similarity between its CORAL-derived spectrum and the corresponding external spectrum.

In the Multiz100 comparison (Figure S2a), human and chimpanzee (*P. troglodytes*) showed nearly identical spectra (cosine = 0.9995 and 0.9991, respectively). Across species, cosine similarity between matching spectra showed a significant negative correlation with evolutionary distance from humans (Spearman = -0.564, p = 0.00267) and no positive correlation with alignment coverage (Spearman = -0.316, p = 0.115). These findings highlight the value of the non-single-reference alignment strategy and indicate that differences in alignment coverage do not systematically bias the inferred spectra.

We next compared CORAL spectra with polymorphism-based mutation spectra reported in *msad213* for 13 species ^7^ (Figure S2b). For matching species, nearly all pairs showed cosine similarity greater than 0.98, and matching pairs were significantly more similar than non-matching pairs (mean 0.988 vs. 0.954; one-sided Wilcoxon rank-sum test, p = 1.69×10^-6^). In addition, species’ spectra were consistently more similar to those of closely related taxa than to distantly related ones, motivating further exploration of the phylogenetic structure embedded in mutation spectra.

To further validate the accuracy of CORAL spectra, we compared fungal mutation spectra inferred by CORAL with spectra derived from three-way Multiz ^8^ alignments across 65 fungal triplets, yielding 130 fungal species-branches. Importantly, fungi exhibited low pseudo-read alignment fractions, making them a particularly challenging clade. For each species-branch, spectra were compared using cosine similarity, yielding a CORAL-by-Multiz similarity matrix (Figure S2c). Matched similarities were extremely high (diagonal mean = 0.9944; median = 0.9952) and markedly exceeded non-matched pairs (off-diagonal mean = 0.9178; median = 0.9466). This diagonal enrichment was highly significant (one-sided Mann–Whitney U test, p = 2.47×10^-82^), indicating that CORAL recapitulates branch-specific fungal spectra with near-identical composition to Multiz in alignable regions.

However, mutation rates showed a different relationship with alignment coverage. Comparison with Zoonomia ^9^ mutation rates showed that low alignment coverage biases mutation-rate estimates (Figure 2a), whereas mutation spectra remained stable across coverage levels (Figure S2; Figure S7c). To avoid including potentially unreliable branches, analyses that depend directly on mutation counts, such as those in comparison of traits with mutation rates, and the distribution of mutation rate across clades (Figure 2c,d), were restricted to species-branches with more than 25% aligned reads. We applied the same threshold to de novo signature extraction as a precaution, ensuring that only well-supported branches were used for calculation.

For all other analyses based on mutation spectra and their derived features, all species-branches were included. Notably, all triplets were retained when computing in-triplet features (e.g., sister-taxa mutation ratios), as mutations are called symmetrically from outgroup-aligned regions and are therefore less sensitive to absolute rate biases.

**Figure S2.**
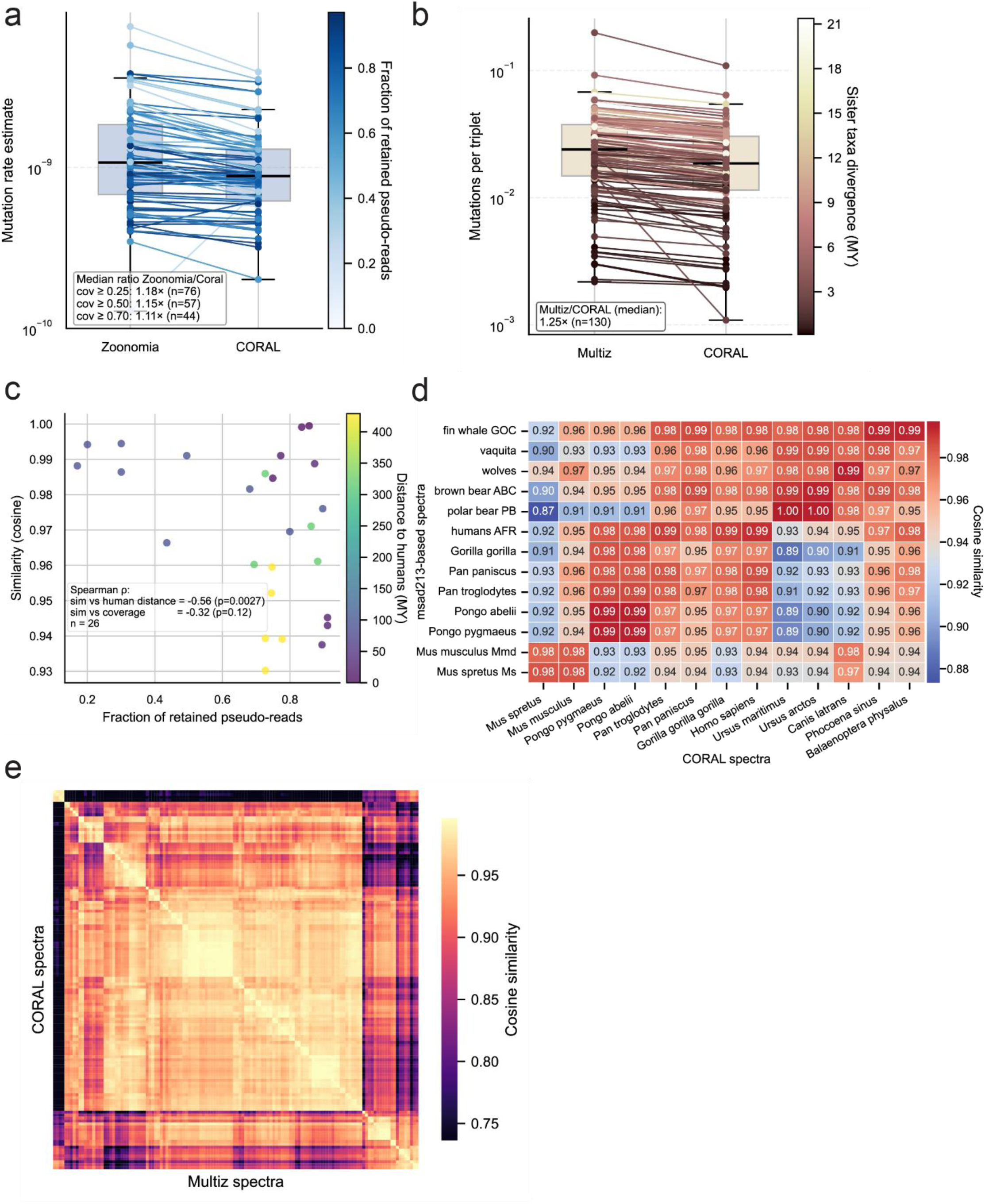
External validation of CORAL-derived mutation rates and spectra. **a,** Paired boxplot comparing CORAL- and Zoonomia-derived mutation-rate estimates (right and left boxplots, respectively). Each dot represents a species-branch estimate, with lines connecting the same branches. Points are colored by the fraction of aligned pseudo-reads retained for that branch in CORAL. The median Zoonomia/CORAL ratio is reported for coverage thresholds of ≥0.25, ≥0.5, and ≥0.7. **b,** Paired boxplot comparing Multiz- and CORAL-derived mutations per callable site for fungal species (left and right boxplots, respectively). Each dot represents a species-branch estimate, with lines connecting the same branches. Points are colored by sister-taxon divergence time, and the annotated value indicates the median Multiz/CORAL ratio. **c,** Cosine similarity between CORAL-derived mutation spectra and spectra extracted from Multiz100 with analogous three-way comparisons, plotted against the fraction of retained pseudo-reads. Each point represents a species and is colored by evolutionary distance from humans (MY). Spearman correlation coefficients and p-values are shown for similarity versus distance from humans and similarity versus alignment coverage. **d,** Cosine similarity heatmap comparing CORAL-derived mutation spectra (columns) with msad213 spectra (rows). Cell values indicate cosine similarity. **e,** Cosine similarity heatmap comparing CORAL-derived mutation spectra (rows) with Multiz-based spectra (columns) for the chosen fungi species. Cell values indicate cosine similarity.

### Supplementary Note 3

#### Time dependence of mutation rates with divergence time

We observed a strong and consistent negative correlation between mutation rate estimates and both sister taxa and outgroup divergence times (Figure 2d; Figure S3b; Figure S5). One possible explanation is coverage bias, arising from the preferential alignment of conserved regions. Indeed, divergence time negatively correlates with alignment coverage (Figure S1f; Figure S3b), which in turn correlates with mutation-rate estimates in some groups. However, even when comparing species with similar alignment coverage, we observed the same time-dependent decline in mutation rate, suggesting that coverage bias is not the primary cause of this pattern (Figure S5b).

Alternative explanations include: mutation saturation, potentially exacerbated by regional clustering of mutations and leading to masking of multiple hits or reduced apparent mutation probability; purifying selection, which gradually removes non-fixed variants over time; and systematic underestimation of divergence times, which would bias mutation-rate estimates upward ^10,11^. Overall, our results reveal a clear hyperbolic time-dependence of mutation rates across phyla (Figure S3c). Reptiles represent a notable exception, showing little or no such trend. This anomaly may reflect their generally low mutation rates, potentially indicating saturation effects or other factors, as reptiles were also outliers in read-alignment behavior (Supplementary Note 1).

**Figure S3.**
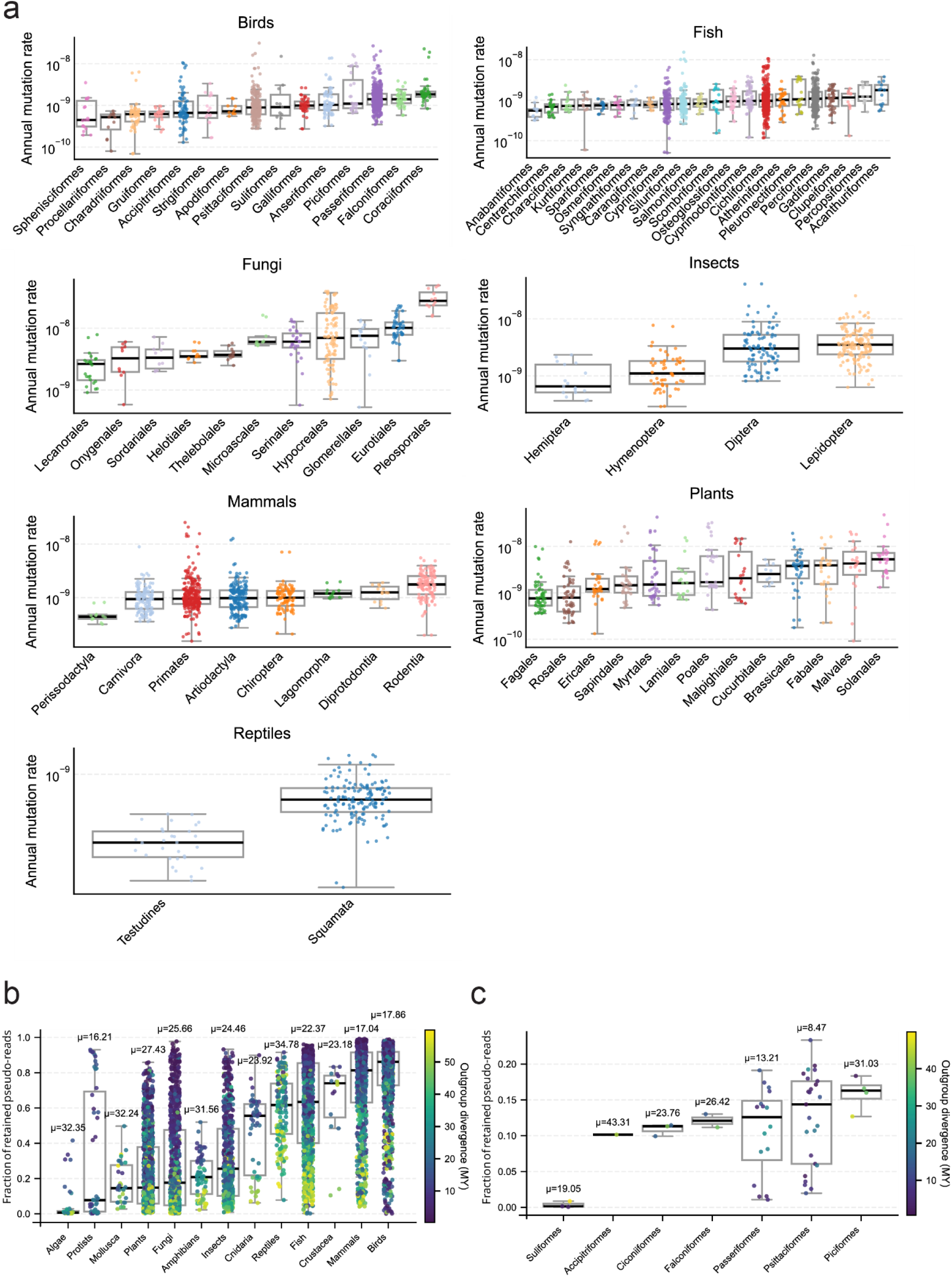
Variation in mutation rates and alignment across taxonomic groups. **a,** Annual mutation rate estimates across orders within each clade (panels). Boxplots summarize per-order distributions; points represent individual species-branches. Only orders with more than 10 species-branches are shown. **b,** Fraction of retained pseudo-reads across clades. Boxplots summarize per-clade distributions; points represent species-branches and are colored by outgroup divergence time (MY). Mean outgroup divergence time (𝜇) is shown above each box. **c,** Fraction of retained pseudo-reads across bird orders with <0.25 fraction of kept pseudo-reads, plotted as in panel **b**.

**Figure S4.**
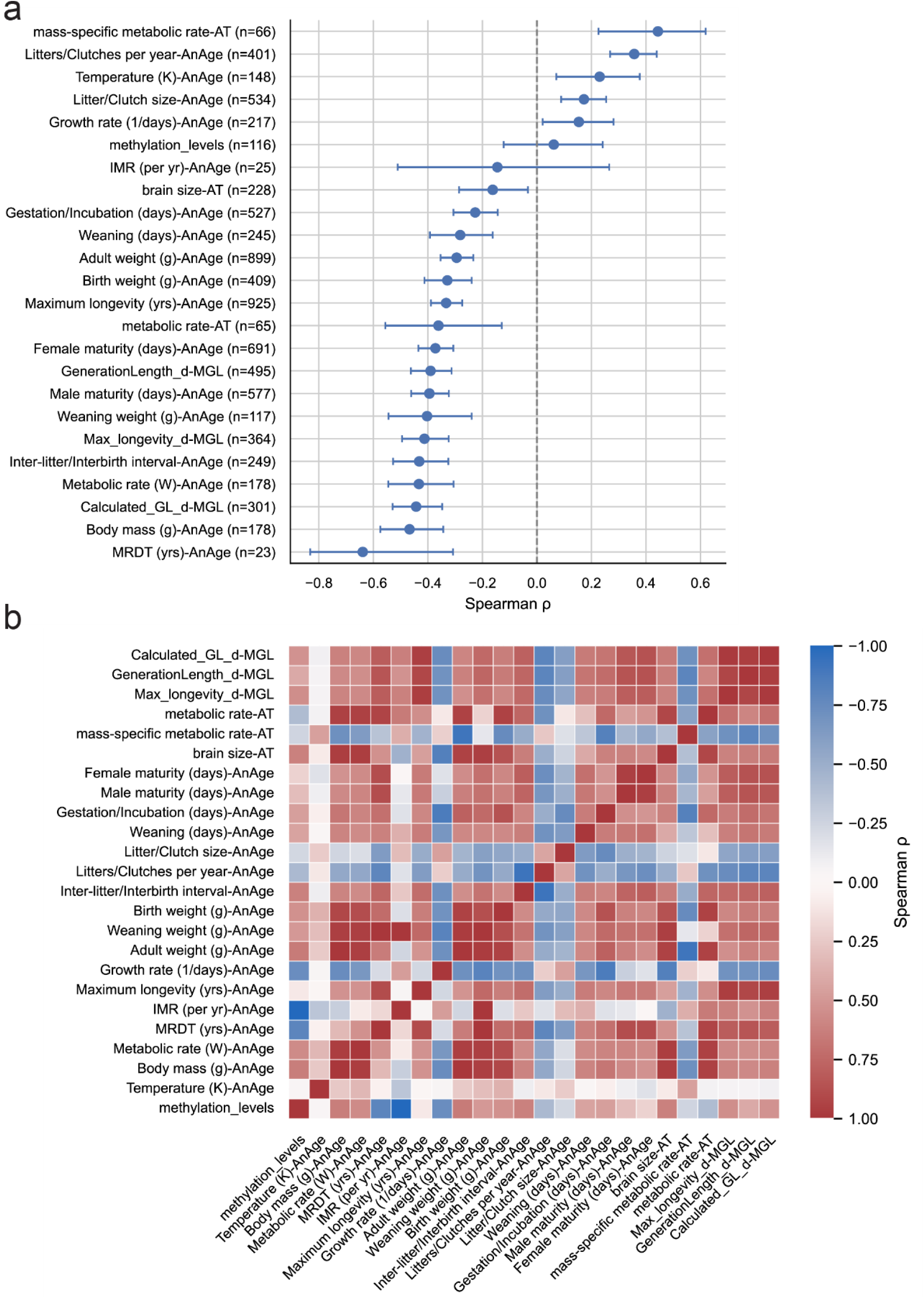
Association of mutation rates with life-history and ecological traits. **a,** Spearman correlation coefficients between mean annual mutation rate per species and life-history traits from AnAge, AnimalTraits (AT), and Pacifici *et al.* Points indicate correlation estimates with 95% confidence intervals; the dashed line denotes zero correlation. P-values are shown in Table S4. **b,** Pairwise Spearman correlation matrix among life-history traits used in the analysis. Cell color indicates correlation strength and direction.

**Figure S5.**
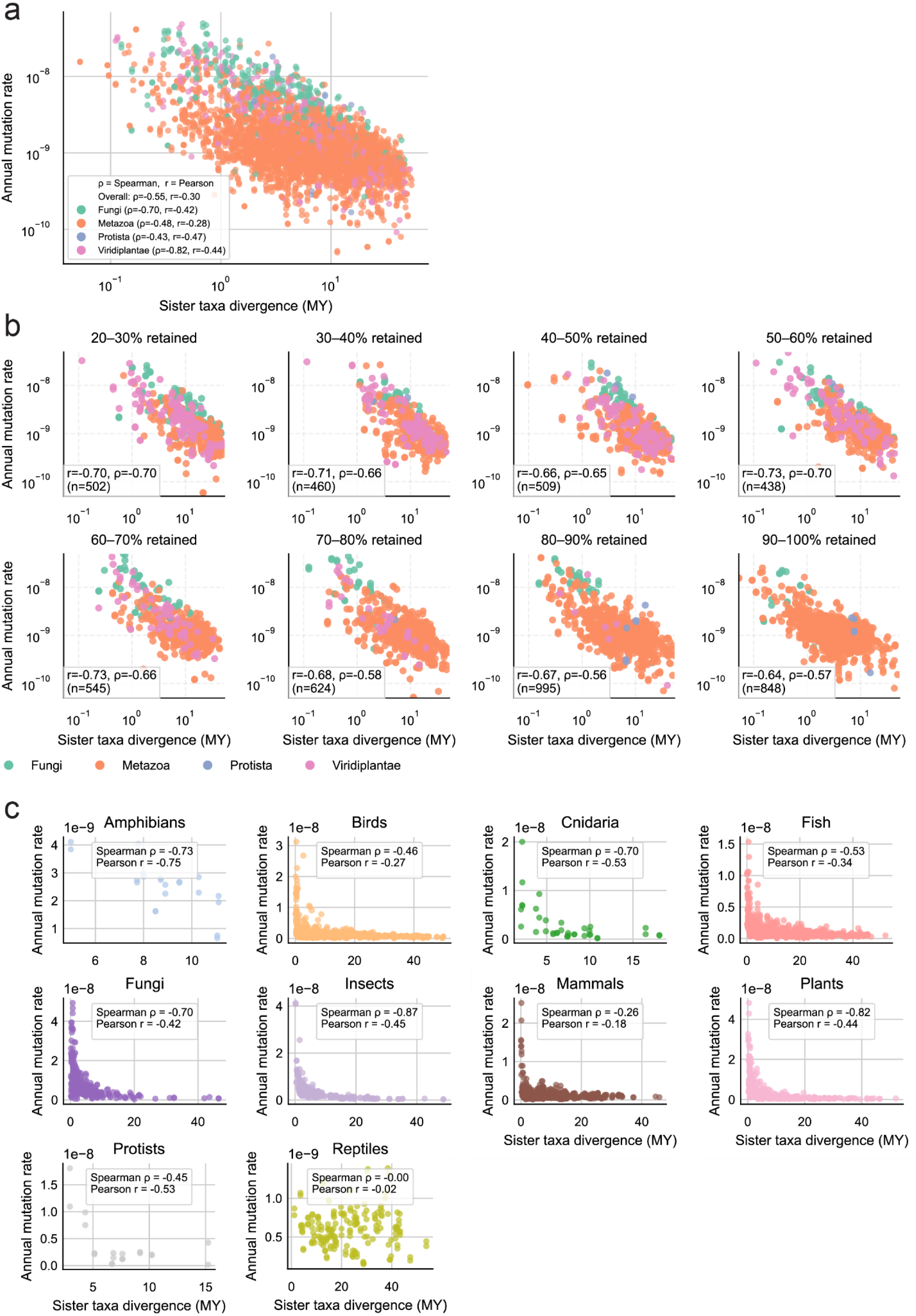
Dependence of mutation rate estimates on divergence time. **a,** Annual mutation rate estimates versus sister-taxa divergence time (MY) across species. Points are colored by kingdom; Spearman and Pearson correlation coefficients are shown overall and per kingdom. b, Annual mutation rate versus sister-taxa divergence time stratified by alignment coverage. Panels show species-branches as points; Spearman correlation coefficients are reported within each coverage bin. **c,** Annual mutation rate versus sister-taxa divergence time shown separately by clade. Each panel corresponds to one clade; points represent species, with Spearman and Pearson correlation coefficients reported.

**Figure S6.**
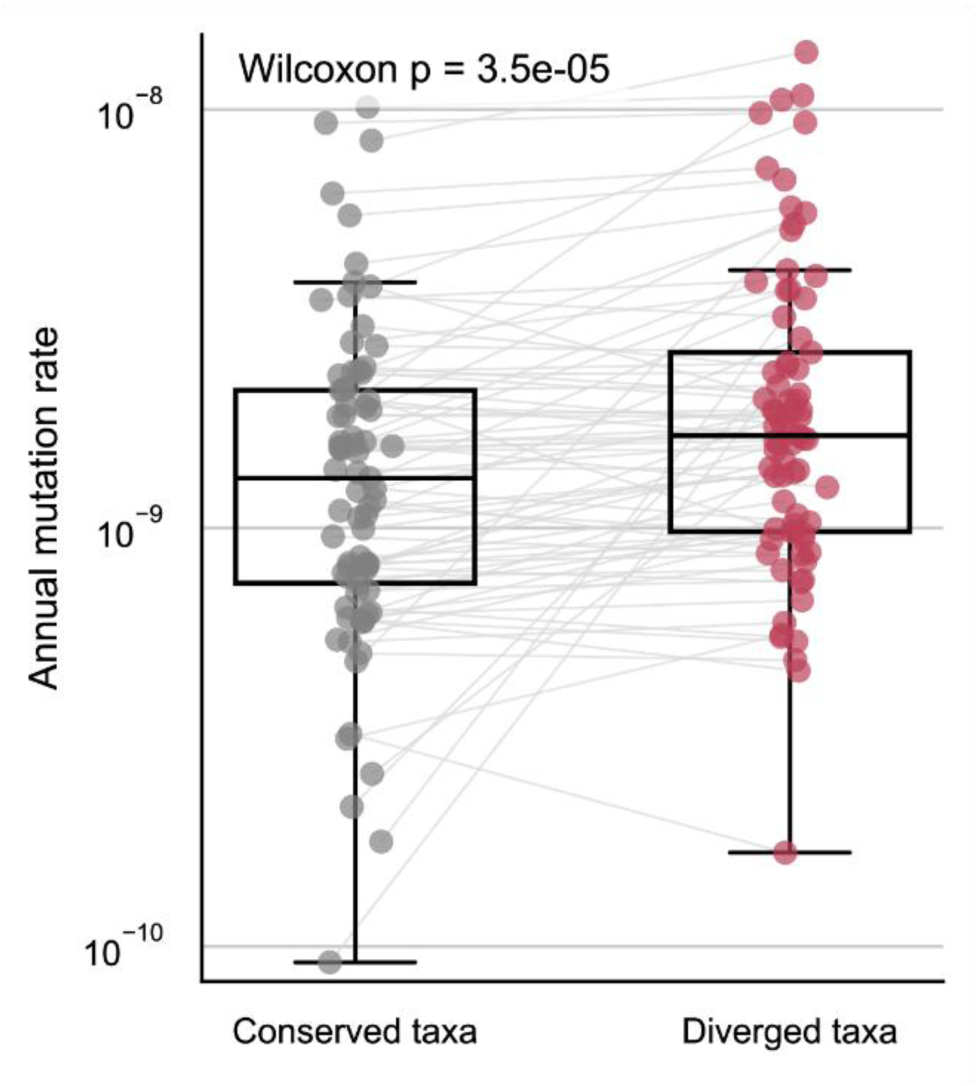
Genus divergence analysis. Annual mutation rates for same-genus and genus-diverged sister taxa within triplets. Sister-taxon estimates are connected by lines; boxplots summarize distributions. The Wilcoxon signed-rank test p-value is shown.

**Figure S7.**
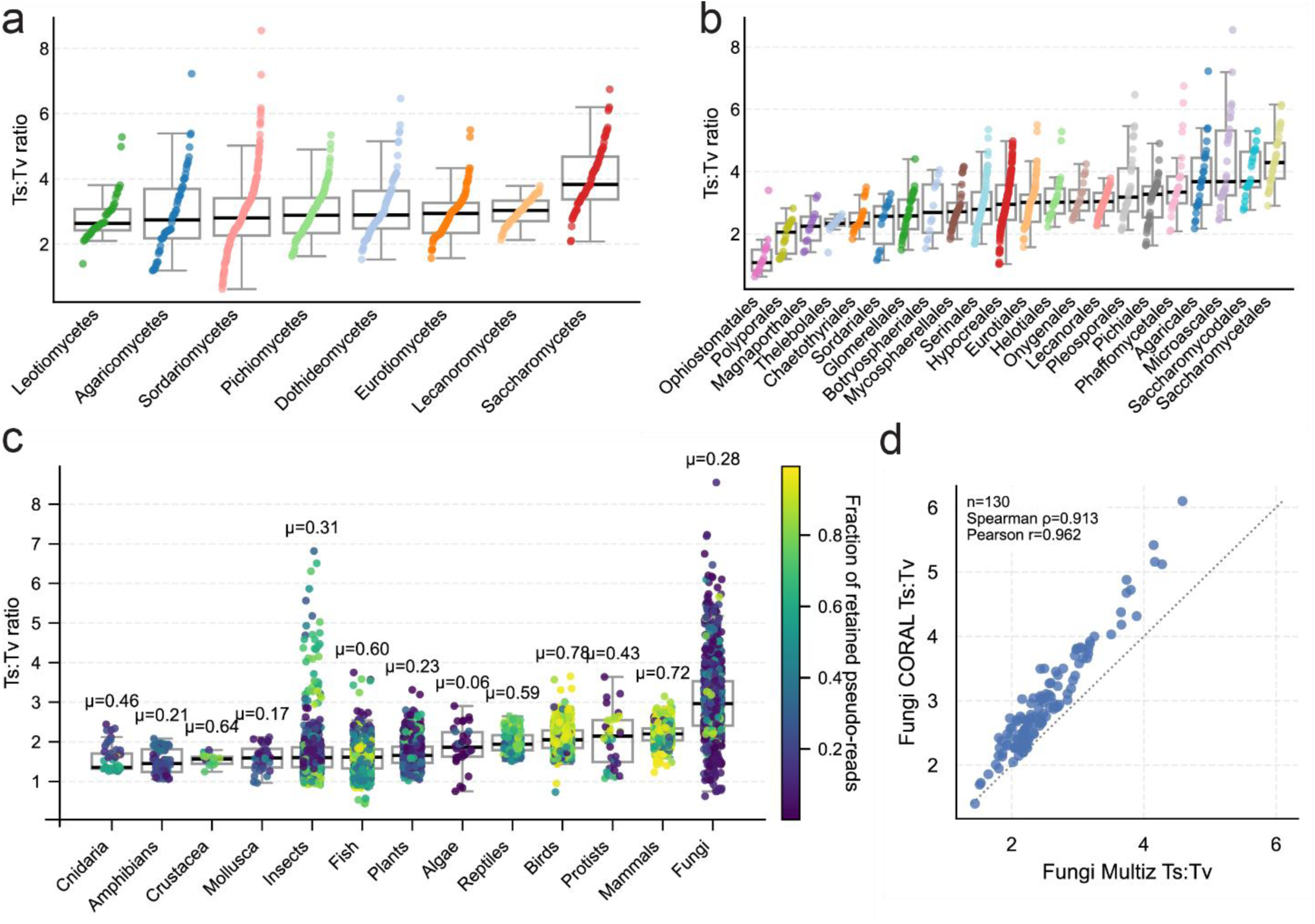
Transition:transversion ratios across taxa. **a,** Transition–transversion (Ts:Tv) ratios across fungal classes. Boxplots summarize class-level distributions; points represent individual species-branches. **b,** Ts:Tv ratios across fungal orders, plotted as in **a**. **c,** Ts:Tv ratios across clades. Boxplots summarize per-clade distributions; points represent species-branches and are colored by the fraction of retained pseudo-reads. The mean fraction of retained pseudo-reads (μ) is indicated above each clade. **d,** Comparison of Ts:Tv ratios between CORAL (y-axis) and Multiz (x-axis) for selected fungal species-branches. Spearman and Pearson correlation coefficients are shown.

**Figure S8.**
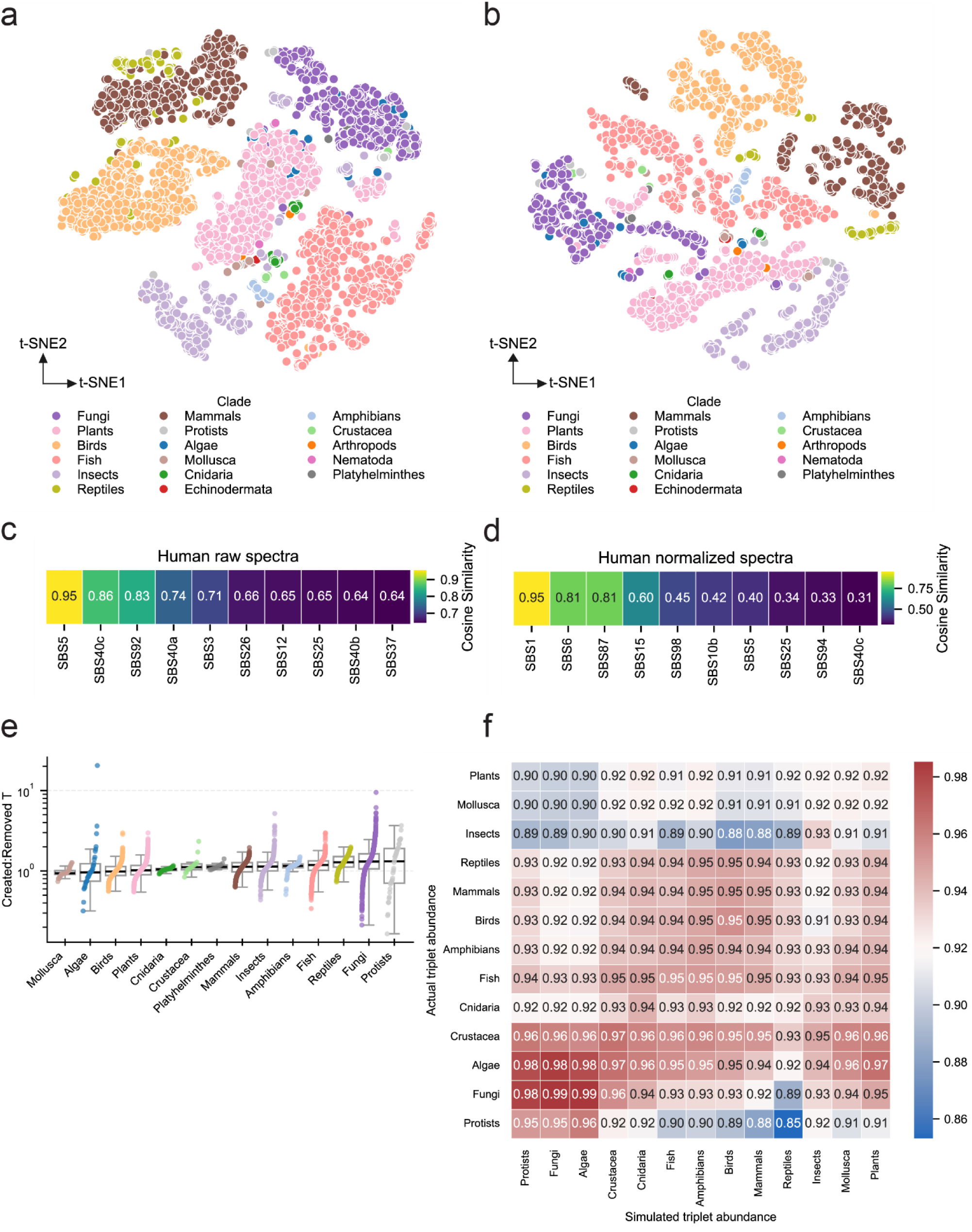
Normalization of mutation spectra and relationships to trinucleotide composition. **a,** t-SNE embedding of 96-category pre-normalized mutation spectra across species-branches, with points colored by clade. **b,** t-SNE embedding of 32-category 3-mer abundance across species, plotted as in **a**. **c,** Cosine similarity between the raw human mutation spectrum and top-most similar COSMIC SBS signatures; values indicate cosine similarity. **d,** Cosine similarity between the normalized human mutation spectrum and top-most similar COSMIC SBS signatures, plotted as in **c**. **e,** Ratios of created to removed AT bases across clades (Methods). Boxplots summarize per-clade distributions; points represent species-branches. **f,** Cosine similarity matrix comparing observed trinucleotide abundances (rows) with steady-state compositions simulated from clade-mean normalized mutation spectra (columns). Values indicate cosine similarity.

**Figure S9.**
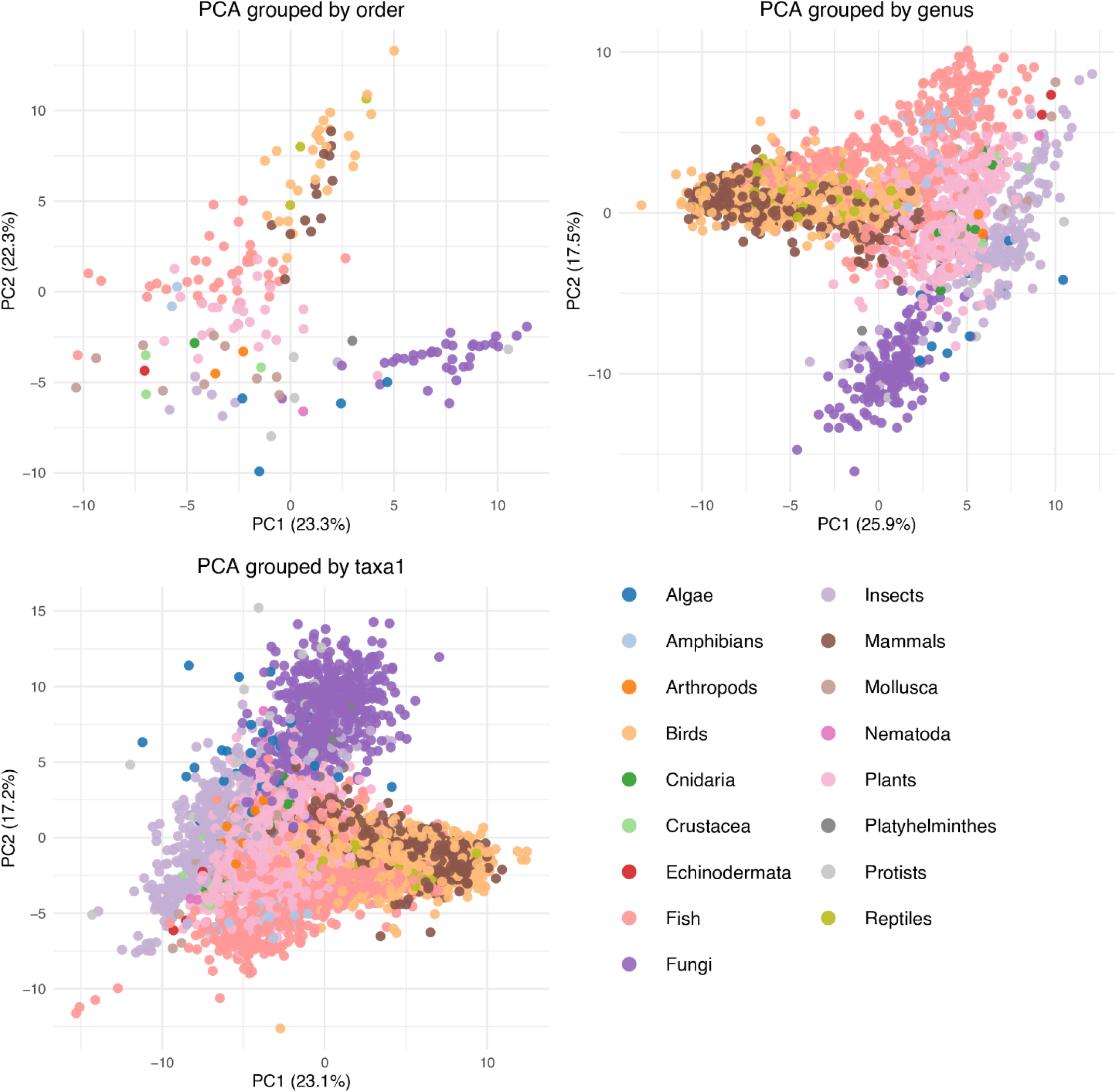
Principal component analysis of normalized mutation spectra. PCA plots showing the first two principal components of mean normalized mutation spectra at three taxonomic resolutions: orders (top left), genera (top right), and species (bottom left). Points represent groups of species-branches at the corresponding level and are colored by clade. Axes indicate the variance explained by each principal component.

**Figure S10.**
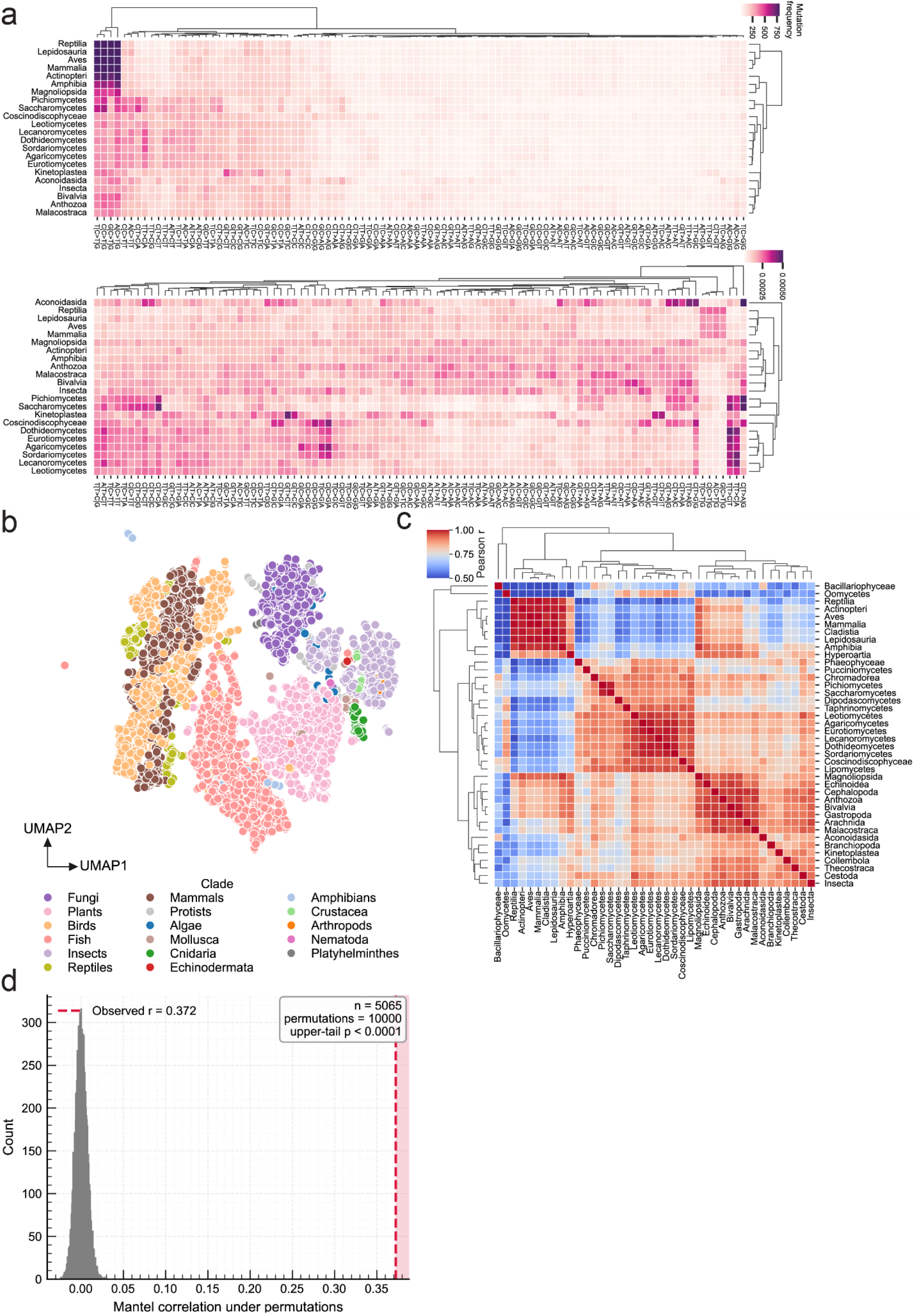
Phylogenetic structure of normalized mutation spectra. **a,** Hierarchical clustering of clade-level mutation spectra based on pairwise spectral similarity. Rows correspond to clades and columns to mutation categories. The top heatmap shows mean normalized mutation spectra per clade. The bottom heatmap shows the same spectra further normalized across clades so that each mutation category sums to 1, highlighting relative differences among clades. **b,** UMAP embedding of normalized mutation spectra across species-branches, with points colored by clade. **c,** Pairwise similarity cluster map between mean normalized mutation spectra of classes. Color indicates correlation strength. **d,** Distribution of Mantel test statistics comparing spectral similarity and phylogenetic distance. The observed statistic is indicated relative to the permutation null distribution (n permutations shown).

**Figure S11.**
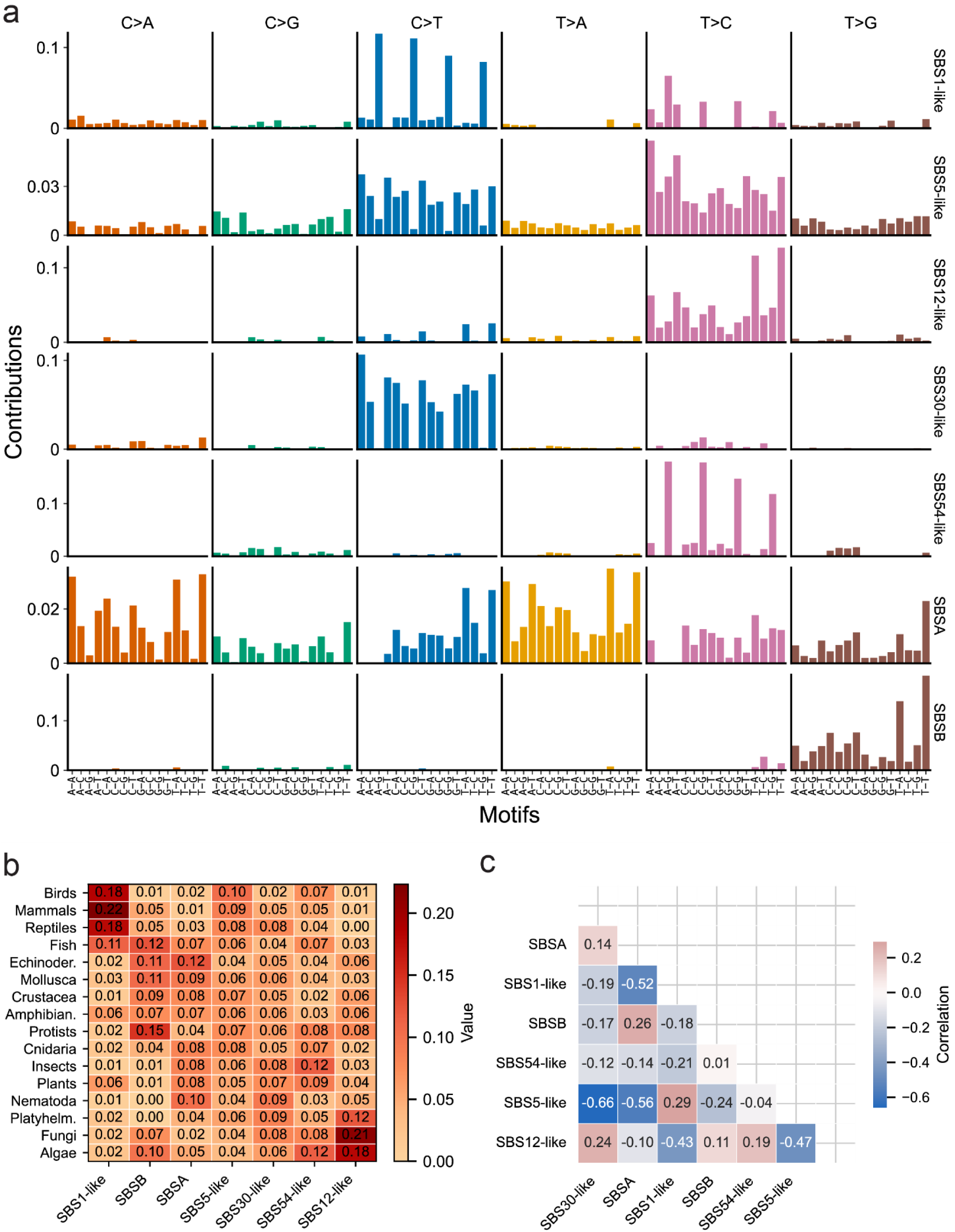
De novo evolutionary mutational signatures. **a,** 96-category mutation spectra of the seven de novo evolutionary signatures identified by unsupervised non-negative matrix factorization (NMF; SignatureAnalyzer). Columns correspond to substitution classes and trinucleotide contexts; rows correspond to signatures. **b,** Relative mean activity of each de novo signature across clades. For each signature, values indicate the fraction of total activity in each clade. **c,** Pairwise correlation matrix of de novo signature activities across species-branches. Cell values indicate correlation coefficients.

**Figure S12.**
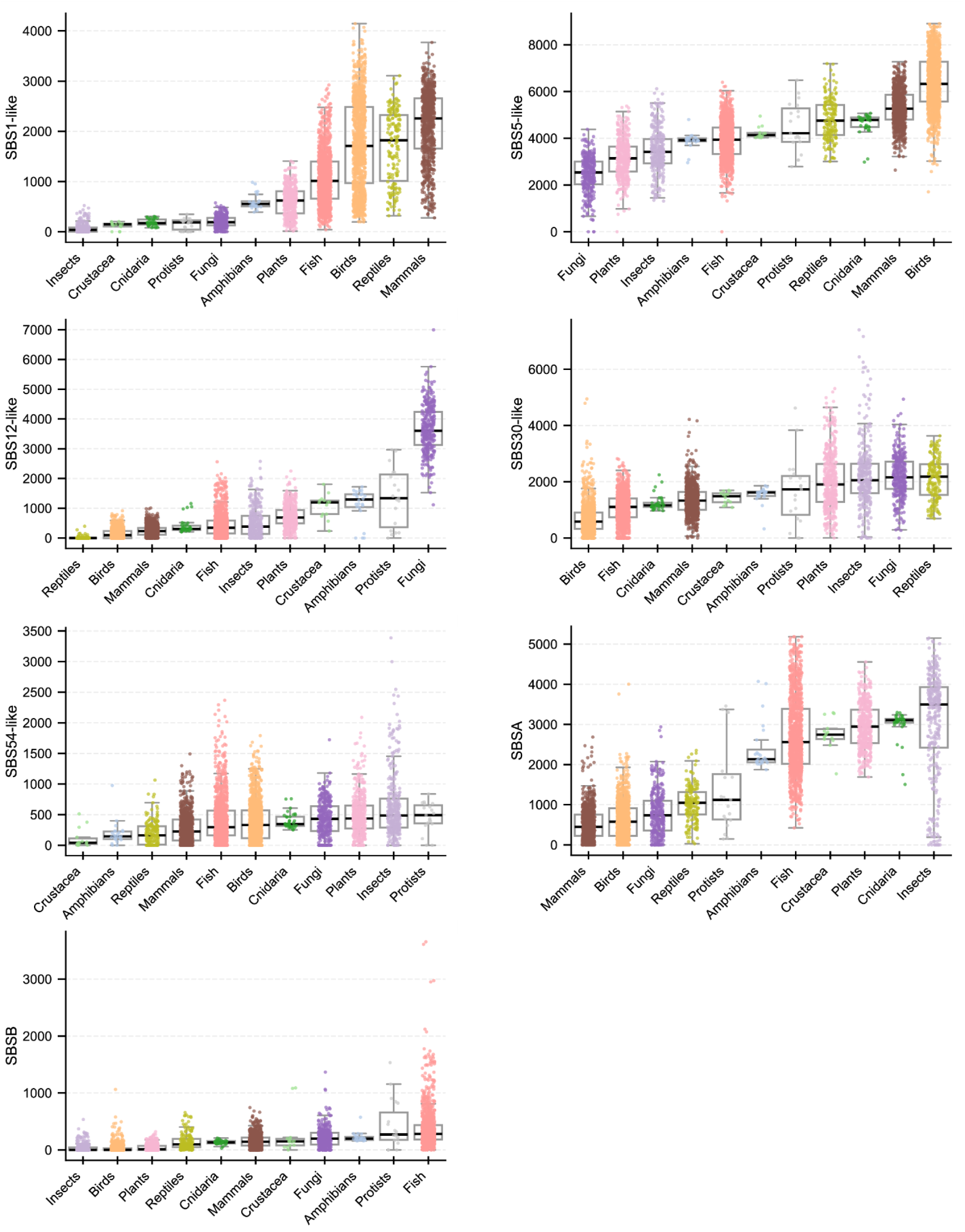
De novo mutational signature activity. Distribution of de novo signature activities across clades. Each panel corresponds to one signature; points represent species-branches and boxplots summarize per-clade distributions.

**Figure S13.**
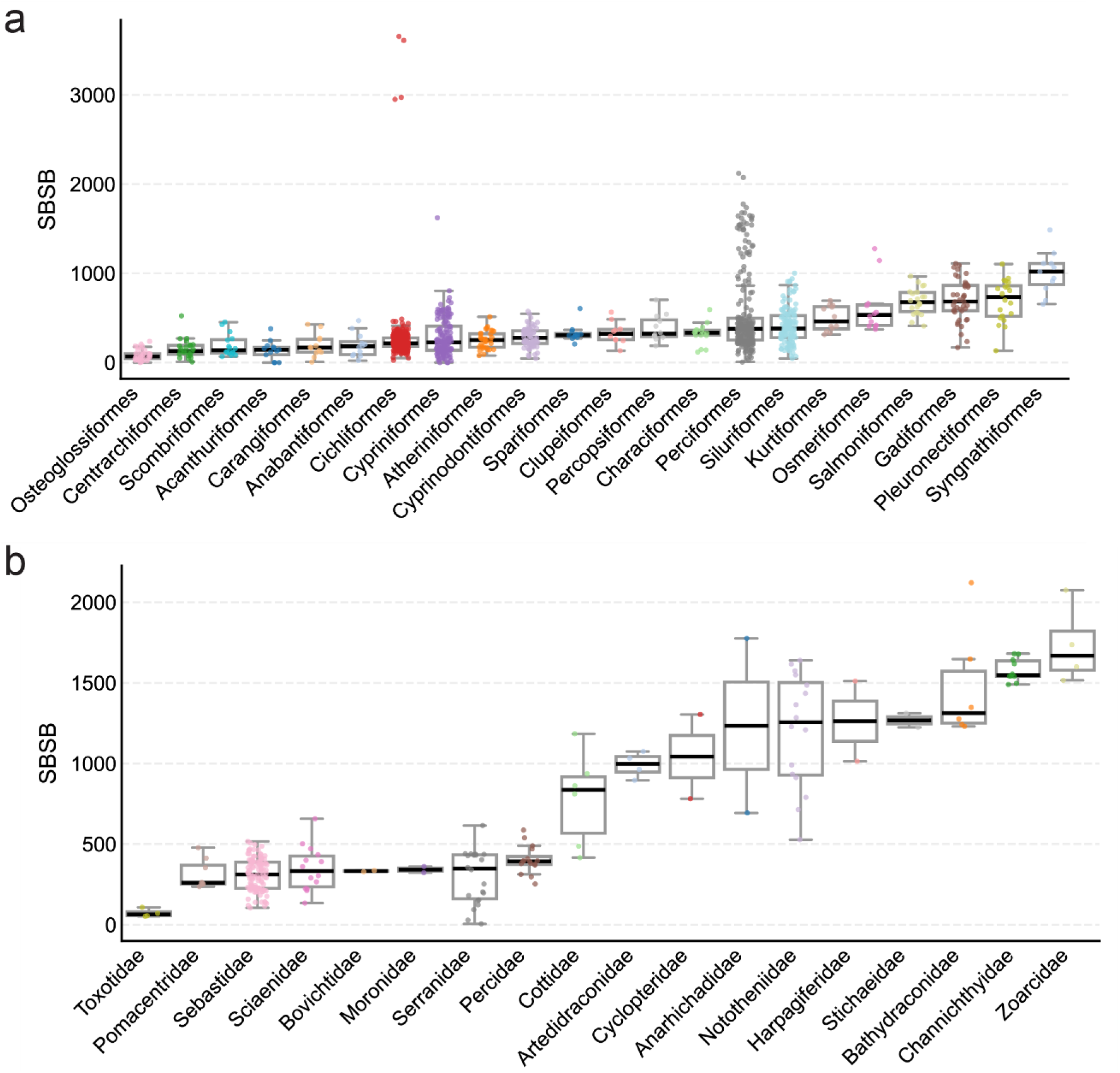
SBSB signature activity. **a,** Distribution of SBSB activity across fish orders. Points represent species-branches, and boxplots summarize per-order distributions. **b,** Similar to **a.** across Perciformes fish families.

**Figure S14.**
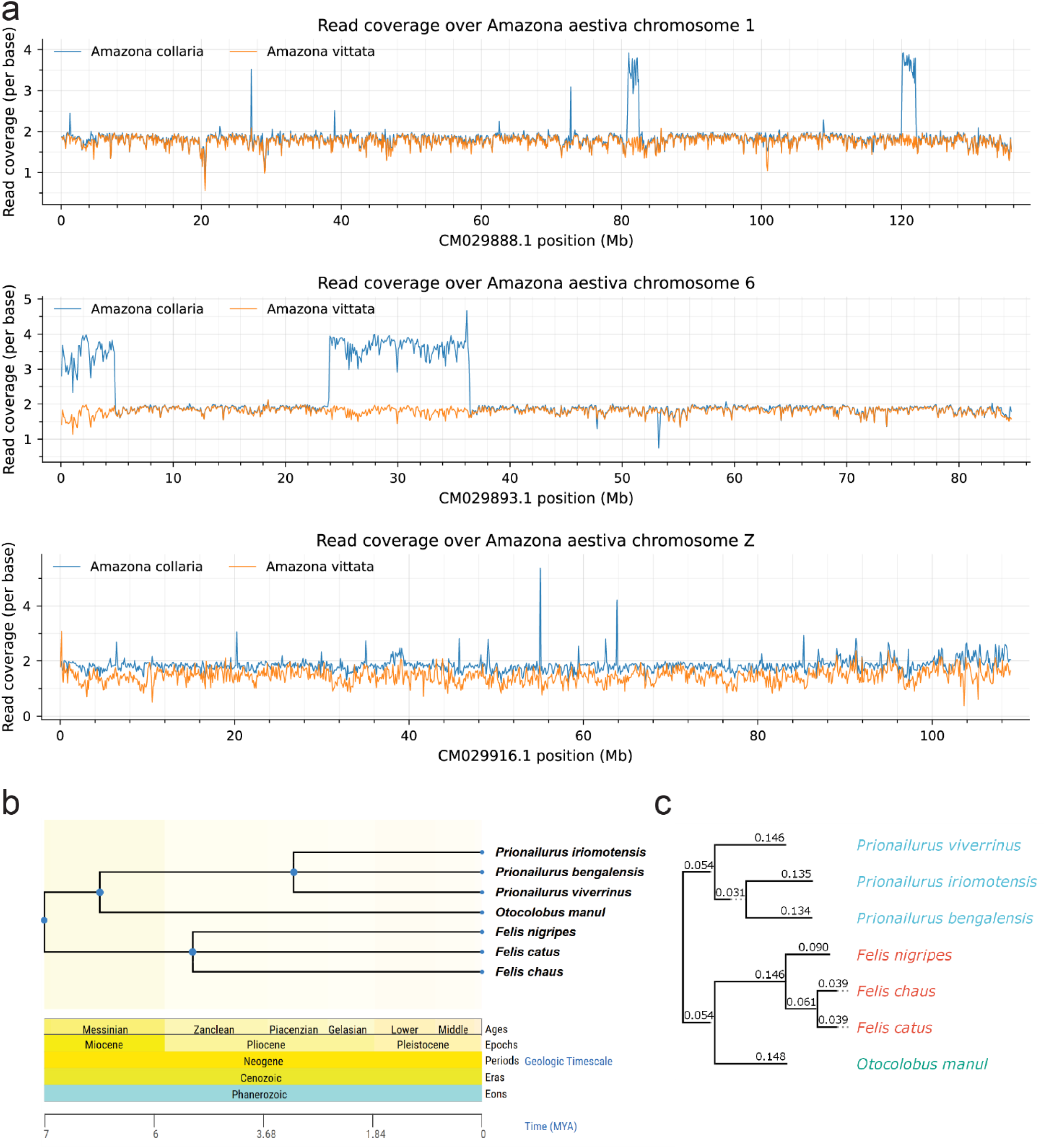
CORAL extended applications. **a,** Read coverage profiles for *Amazona collaria* and *A. vittata*, aligned to the A. aestiva reference genome across reference chromosomes 1 (top), 6 (middle), and Z (bottom). Smoothed read depth is shown along genomic position (100,000bp sliding window). **b,** Reference phylogenetic relationships among selected Felidae species based on TimeTree v5. Branch lengths reflect divergence-time estimates. **c,** Parsimony-based phylogenetic reconstruction inferred from CORAL multi-species alignment. Pseudo-reads from all species were aligned to *Otocolobus manul* as a common reference, and variable sites differing in at least one species were extracted and used for tree reconstruction with PHYLIP. Branch lengths reflect relative substitution number.

### Supplementary Note 4

#### Genomic position and coverage analysis

Beyond genome-wide spectra, CORAL extracts positional information on read coverage and mutation density along the outgroup genome, enabling detection of local alterations in alignment depth and mutation distribution. Coverage is derived from the raw BAM files for each sister taxon and summarized along the outgroup genome using a sliding-window approach (100,000 bp), which facilitates direct comparison of coverage profiles between aligned species and highlights large-scale copy number and deletion events.

We demonstrate this potential using a triplet of *Amazona* species. Aligning *A. vittata* and *A. collaria* to *A. aestiva* (outgroup) revealed several notable patterns (Figure S14a). First, read coverage across the genome was close to the expected 2×, and local fluctuations in coverage were largely shared between sister taxa, reflecting either shared evolutionary changes or missing segments in the reference genome. Nonetheless, certain regions showed a doubling to nearly 4× in *A. collaria*, particularly on chromosome 1 (80–83 Mb and 120–122 Mb) and chromosome 6 (0–5 Mb and 24–36 Mb). The consistency of this doubling strongly suggests true biological duplications. This illustrates how read-based approaches similar to those used in cancer genomics can be applied in a comparative evolutionary context in order to identify large copy-number variants.

Examining the *Amazona* Z chromosome revealed a markedly different pattern. Unlike the autosomes, where both sister taxa exhibited nearly identical coverage, coverage along chromosome Z diverged substantially between species. This suggests more rapid sequence evolution and lineage-specific changes, consistent with accelerated sex-chromosome evolution driven by sex-biased inheritance ^12^, and the atypical sex-chromosome evolution reported in parrots ^6^.

Interpretation of such patterns warrants caution, as sequencing biases and differences in repeat handling may affect alignment. Nevertheless, the clear regional coverage changes observed here underscore the potential of this approach to detect large-scale genomic rearrangements across the tree of life.

### Multi-species branch-wise phylogenetic detection

Although this work focused on pairwise sister-taxon comparisons, the same framework can be extended to multi-species alignments against a common reference. Aligning more than two species enables direct comparison of orthologous sites across taxa, broadening inference from mutations along terminal branches to ancestral branches within the phylogenetic tree, and supporting fine-scale phylogenetic reconstruction of recent divergence history.

As a proof of concept, we examined several cat species (*Otocolobus manul*, *Felis catus*, *Felis chaus*, *Felis nigripes*, *Prionailurus bengalensis*, *P. iriomotensis*, and *P. viverrinus*). Their evolutionary relationships remain partly unresolved in *TimeTree5* (TTol5) ^13^ due to limited sequence data (Figure S14b). Aligning all species’ pseudo-reads to *O. manul* as the reference, we extracted variable sites differing in at least one species and used these sites for phylogenetic reconstruction with PHYLIP ^14^. This reconstruction resolved two ambiguous nodes: within Felis, F. nigripes diverged first, while F. chaus and F. catus formed a sister pair (Figure S14c). Within Prionailurus, P. bengalensis and P. iriomotensis appeared as sister taxa. Notably, a previous study proposed *F. nigripes* and *F. catus* as sister taxa, in contrast to our finding ^15^. Although preliminary and requiring more stringent site-level filtering, this example illustrates how CORAL outputs can inform phylogenetic reconstruction, particularly among closely related species.

## Supplementary table legends

**Supplementary Table 1 Lean CORAL database**

Per-branch mutation data derived from species triplets. Each row represents a focal lineage from a sister-outgroup triplet and reports raw and normalized 96-category mutation counts, callable trinucleotide context abundances, taxonomy and divergence times, alignment and mutation-rate metrics, collapsed substitution summaries, de novo mutational signature activities, and external benchmarks and life-history traits.

**Supplementary Table 2 CORAL alignment statistics**

Pseudo-read alignment and filtering statistics for all species-branches (Sheet 1), including retained read fractions and filtering causes, and alignment runtime and genome size summaries for each species triplet (Sheet 2).

**Supplementary Table 3 Taxonomic coverage summary**

Counts of species-branches and unique species represented in the CORAL dataset, summarized by kingdom, phylum, and class, reported both before and after applying a ≥25% alignment coverage threshold.

**Supplementary Table 4 Trait correlations with mutation rate**

Spearman correlations between annual mutation rate and life-history traits computed on species-level means, reported for all species combined and separately for each clade.

**Supplementary Table 5 Partial rank correlation (PRCC) analysis**

Partial rank correlation coefficients between mutation rate and life-history traits, controlling for correlated predictors within predefined trait groups (G1-G4; each summarized is a separate sheet), reported for all species and for individual clades.

**Supplementary Table 6 De novo mutational signatures**

De novo mutational signatures inferred by signature analyzer, including signature profiles across 96 mutation channels (Sheet 1), per-branch signature activities (Sheet 2), and cosine similarity to COSMIC SBS signatures (Sheet 3).

**Supplementary Table 7 Signature accumulation with divergence time**

Correlations between total mutation burden or de novo signature burden and sister-taxa divergence time, reported for all species and by clade, using both Pearson and Spearman statistics.

**Supplementary Table 8 Reference genomes and triplet definitions**

NCBI reference genome assemblies considered in this study (Sheet 1) and the final set of species triplets used for CORAL analyses with associated TimeTree divergence times (Sheet 2).

